# The ribose methylation enzyme FTSJ1 has a conserved role in neuron morphology and learning performance

**DOI:** 10.1101/2021.02.06.430044

**Authors:** Mira Brazane, Dilyana G Dimitrova, Julien Pigeon, Chiara Paolantoni, Tao Ye, Virginie Marchand, Bruno Da Silva, Elise Schaefer, Margarita T Angelova, Zornitza Stark, Martin Delatycki, Tracy Dudding-Byth, Jozef Gecz, Pierre-Yves Placais, Laure Teysset, Thomas Preat, Amélie Piton, Bassem A. Hassan, Jean-Yves Roignant, Yuri Motorin, Clément Carré

## Abstract

FTSJ1 is a conserved human 2’-O-methyltransferase (Nm-MTase) that modifies several transfer RNAs (tRNAs) at position 32 and the wobble position 34 in the AntiCodon Loop (ACL). Its loss of function has been linked to Non-Syndromic X-Linked Intellectual Disability (NSXLID), and more recently to cancers. However, the molecular mechanisms underlying these pathologies are currently unclear. Here we report a novel *FTSJ1* pathogenic variant from a NSXLID patient. Using blood cells derived from this patient and other affected individuals carrying *FTSJ1* mutations, we performed an unbiased and comprehensive RiboMethSeq analysis to map the ribose methylation (Nm) on all human tRNAs and identify novel targets. In addition, we performed a transcriptome analysis in these cells and found that several genes previously associated with intellectual disability and cancers were deregulated. We also found changes in the miRNA population that suggest potential cross-regulation of some miRNAs with these key mRNA targets. Finally, we show that differentiation of FTSJ1-depleted human neuronal progenitor cells (NPC) into neurons displays long and thin spine neurites compared to control cells. These defects are also observed in *Drosophila* and are associated with long term memory deficit in this organism. Altogether, our study adds insight into FTSJ1 pathologies in human and flies by the identification of novel FTSJ1 targets and the defect in neuron morphology.

## INTRODUCTION

RNA modifications represent a novel layer of post-transcriptional gene regulation (Saletore *et al*, 2012; Angelova *et al*, 2018; Zhao *et al*, 2020). Due to their variety and dynamic nature, they rapidly adapt gene expression programs in response to developmental changes or environmental variations. One of the most abundant RNA modifications is 2’-O-methylation (ribose methylation, Nm). Nm can affect the properties of RNA molecules in multiple ways *e.g*. stability, interactions and functions (Kawai *et al*, 1992; Kurth & Mochizuki, 2009; Lacoux *et al*, 2012). Nm residues are abundant in ribosomal RNAs (rRNAs) and transfer RNAs (tRNAs) (Erales *et al*, 2017; Marchand *et al*, 2017), but are also found in other RNA types such as small nuclear RNAs (snRNAs) (Darzacq, 2002; Dai *et al*, 2017), small non-coding RNAs (sncRNAs) (Li *et al*, 2005; Yu *et al*, 2005; Horwich *et al*, 2007; Saito *et al*, 2007; Kurth & Mochizuki, 2009) and messenger RNAs (mRNAs) (Darzacq, 2002; Dai *et al*, 2017; Bartoli et al. 2018). Many Nm positions are conserved through evolution and their presence is essential for maintaining healthy physiological functions. Eukaryotic mRNAs are 5’ end capped with a 7-methylguanosine (m^7^G), which is important for processing and translation of mRNAs. In addition, Cap methyltransferases (CMTR) catalyse Nm of the first and second transcribed nucleotides and were shown to be important for innate immune surveillance, neuronal development and activity (Lee *et al*, 2020; Haussmann *et al*, 2022). The loss of certain Nm modifications and/or Nm-modifying enzymes has been associated to various pathological conditions (reviewed in (Dimitrova *et al*, 2019)), including cancers (Liu *et al*, 2017; El Hassouni *et al*, 2019; He *et al*, 2020; Marcel *et al*, 2020) and brain diseases (Jia *et al*, 2012; Abe *et al*, 2014; Guy *et al*, 2015; Cavaillé, 2017).

FTSJ1 is a human tRNA 2’-O-methyltransferase (Nm-MTase), which belongs to the large phylogenetically conserved superfamily of Rrmj/fibrillarin RNA methyltransferases (Bügl *et al*, 2000; Feder *et al*, 2003). Human males individuals bearing a hemizygous loss of function variant in the *FTSJ1* gene suffer from significant limitations both in intellectual functioning and in adaptive behaviour (Froyen *et al*, 2007; Freude *et al*, 2004; Guy *et al*, 2015). Similar phenotypes, including impaired learning and memory capacity, were recently observed in *Ftsj1* KO mice that also present a reduced body weight and bone mass, as well as altered energy metabolism (Jensen *et al*, 2019; Nagayoshi *et al*, 2021). In flies, we recently showed that the loss of the two *FTSJ1* homologs (*i.e* Trm7_32 and Trm7_34) provokes reduced lifespan and body weight, and affects RNAi antiviral defences and locomotion (Angelova and Dimitrova et al. 2020). Finally, *Ftsj1* mutants in yeast (Δ*trm7*) grow poorly due to a constitutive general amino acid control (GAAC) activation and the possible reduced availability of aminoacylated tRNA^Phe^ (Pintard *et al*, 2002; Guy *et al*, 2012a; Han *et al*, 2018). Interestingly, this growth phenotype can be rescued by human FTSJ1, indicating a conserved evolutionary function.

Most of the knowledge on FTSJ1’s molecular functions are derived from yeast studies. Trm7 in *Saccharomyces cerevisiae* methylates positions 32 and 34 in the AntiCodon Loop (ACL) of specific tRNA targets: tRNA^Phe(GAA)^, tRNA^Trp(CCA)^ and tRNA^Leu(UAA)^ (Pintard *et al*, 2002; Guy *et al*, 2012a). To achieve 2’-O-methylation, Trm7 teams up with two other proteins: Trm732 for the methylation of cytosine at position 32, and with Trm734 for the methylation of cytosine or guanine at position 34 (Guy *et al*, 2012a; Li *et al*, 2020a). The presence of both Cm_32_ and Gm_34_ in tRNA^Phe(GAA)^ is required for efficient conversion of m^1^G_37_ to wybutosine (yW_37_) by other proteins. This molecular circuitry is conserved in the phylogenetically distinct *Schizosaccharomyces pombe* and humans (Noma *et al*, 2006; Guy & Phizicky, 2015; Guy *et al*, 2015; Li *et al*, 2020a). In *Drosophila*, we found that Trm7_32 and Trm7_34 modify, respectively, positions 32 and 34 in the ACL on tRNA^Phe(GAA)^, tRNA^Trp(CCA)^ and tRNA^Leu(CAA)^ (Angelova and Dimitrova et al. 2020). In this organism, we also identified novel tRNA targets for these two enzymes (tRNA^Gln(CUG)^ and tRNA^Glu(CUC)^), which raised the question about their conservation in humans. A recent publication reported that human FTSJ1 modifies position 32 of another tRNA^Gln^ isoacceptor, tRNA^Gln(UUG)^ (Li *et al*, 2020a). This study performed in HEK293T cells tested a selected subset of tRNAs using tRNA purification followed by MS analysis. It was shown that position 32 of tRNA^Arg(UCG)^, tRNA^Arg(CCG)^ and tRNA^Arg(ACG)^ as well as position 34 on tRNA^Arg(CCG)^ and tRNA^Leu(CAG)^ are also 2’-O-methylated by human FTSJ1. tRNA^Arg(ACG)^ was originally identified as a target of fly Trm7_32 (Angelova and Dimitrova et al. 2020), while human tRNA^Leu(CAA) (Kawarada *et al*, 2017)^ and yeast tRNA^Leu(UAA)^ (Guy *et al*, 2012a) were predicted targets of FTSJ1 and Trm7, respectively. However, a comprehensive and unbiased (not selected) analysis of all possible FTSJ1 tRNA targets was not performed, particularly in human patient samples, leaving the full spectrum of FTSJ1 tRNA substrates yet to be identified.

Previously, the enzymatic activity of mammalian FTSJ1 on selected tRNAs has been revealed through HPLC (High-Performance Liquid Chromatography) (Guy *et al*, 2015) and more recently through UPLC-MS/MS (Ultra-Performance Liquid Chromatography–Mass Spectrometry/Mass Spectrometry) (Li *et al*, 2020a; Nagayoshi *et al*, 2021). Both approaches analyse mononucleotides derived from selected tRNAs and are based on already reported sequences. The exact position of the modified nucleotide was thus inferred from available information on tRNA sequences and modification profiles database (Jühling *et al*, 2009; Chan & Lowe, 2016; Boccaletto *et al*, 2018). Recently, a new method called RiboMethSeq was established and allows the identification of Nm sites in a complete unbiased manner, based on the protection conferred by the ribose methylation to alkaline digestion (Marchand *et al*, 2016, 2017). This offers the possibility to identify every Nm site regulated by a particular enzyme, especially when investigating abundant RNA, such as tRNA.

In this study we took advantage of this novel approach to identify the full set of FTSJ1’s tRNA targets in human. We report a novel FTSJ1 pathogenic variant from a NSXLID patient. Using blood cells derived from this affected individual and other individuals carrying distinct *FTSJ1* mutations, we performed an unbiased and comprehensive RiboMethSeq analysis to map the ribose methylation on all tRNAs and reveal new targets. In addition, we performed a transcriptome analysis in these FTSJ1 depleted cells and found that several genes previously associated with intellectual disability (ID) and cancers were deregulated. We also found changes in the miRNA population that suggest potential cross-regulation of some miRNAs with these key mRNA targets. Finally, in accordance with the known importance of FTSJ1 during brain development in mice and its involvement in intellectual disability in humans, we showed that human Neuronal Progenitor Cells (NPC) with inactivated FTSJ1 present abnormal neurite morphology. We also observed this phenotype in *Drosophila* as well as a specific deficit in long term memory. Altogether, our study reveals new targets potentially involved in FTSJ1 pathologies in human and demonstrates a conserved function in neuron morphology and function.

## MATERIALS & METHODS

### FTSJ1 variants and lymphoblastoid cell lines (LCLs)

The various lymphoblastoid cell lines (LCLs) were generated using established methods from blood samples of NSXLID affected or healthy male individuals. The cells were cultured in RPMI-1640 medium with L-glutamine and sodium bicarbonate (ref. R8758-500ML, SIGMA) supplemented with 10% FBS (Gibco) and 1% penicillin–streptomycin (ref. P0781, SIGMA) at 37 °C with 5%CO_2_. Cells were split at ½ dilution approximately 24h before being collected for RNA extraction with TRI-Reagent (Sigma Aldrich) following the manufacturer’s instructions.

#### 6514AW & 6514JW (LCL65AW & LCL65JW in this study)

Family A3 - LCLs from two brothers with mild or severe ID associated with psychiatric manifestations (anger, aggression, anxiety, depression, schizophrenia requiring medication) bearing a splice variant in *FTSJ1*: c.121+1delG (Freude *et al*, 2004). This variant leads to a retention of intron 2, creating a premature stop codon (p.Gly41Valfs*10). Part of the transcripts undergo nonsense-mediated mRNA decay.

#### 11716IJ (LCL11 in this study)

Family A18 - LCL from one male with moderate to severe intellectual disability without dysmorphic features carrying an interstitial microdeletion at Xp11.23. The extent of the deletion was subsequently delineated to about 50 kb by regular PCR and included only the *SLC38A5* and *FTSJ1* genes. qPCR with the FTSJ1-ex3 primers is negative, thus demonstrating the complete deletion of *FTSJ1* locus (Froyen *et al*, 2007).

#### 22341SR (LCL22 in this study)

Family 7 (A26P) - LCL from one male with moderate ID and psychiatric features (mild anxiety and compulsive behavior) carrying a missense mutation c.76G>C; p.Ala26Pro in *FTSJ1*. This family has been reported previously (Guy *et al*, 2015).

#### LCL-MM

This is a newly reported family. The LCL has been generated from one male with mild ID, facial dysmorphia (hypertelorism, pointed chin, ears turned back), speech delay, attention disorders and behavior problems carrying a hemizygous *de novo* variant c.362-2A>T in *FTSJ1*. The mutation is predicted to disrupt the acceptor splice site of exon 6 (NM_012280.3: c.362-2A>T). This variant causes a skipping of the entire exon 6 in the mRNA (r.362_414del) leading to a frameshift and a premature stop codon (p.Val121Glyfs*51) (Figure S1A). Part of the transcripts undergo nonsense-mediated mRNA decay (Figure S1C). Consequently, a strong decrease of the corresponding mRNA steady state level is observed (Figure S1B). This variant was deposited in the *ClinVar* database ([VCV000981372.1](https://www.ncbi.nlm.nih.gov/clinvar/variation/981372/)). The research on LCL-MM was performed according to a research protocol approved by a local Ethics Committee (Comité Consultatif de Protection des Personnes dans la Recherche Biomédicale - CCPPRB). A written informed consent was obtained from the patient and his legal representatives.

<u>18451PK</u> (LCL18 in this study), <u>16806JD</u> (LCL16 in this study), <u>3-2591</u> (LCL25 in this study) and 3-5456 (LCL54 in this study): LCL established from control males. Four LCLs not mutated in the *FTSJ1* gene from unaffected males of similar age were used as controls. A written informed consent was obtained from those individuals and previously described LCLs from patients and their legal representatives in the original publications described above.

### LCL MM variant characterization at the mRNA level

As the FTSJ1 mRNA was highly downregulated in LCL MM, characterization of the FTSJ1 transcript for this experiment was performed on total RNAs from cells treated with cycloheximide (see NMD inhibition protocol below). This allowed a three fold increase in FTSJ1 mRNA in LCL MM (Figure S1B). 1 μg of total RNAs from wild type LCL 25 and LCL MM were treated with DNAse I (M0303S-NEB), and reverse transcription was carried out with random hexamer primers (S0142-Thermo Scientific™) using SuperScript™ III Reverse Transcriptase (18080-044-Invitrogen), following the supplier’s protocol. FTSJ1 cDNAs were amplified from 2μL of RT reaction using the following PCR primers: (Forward: 5’-GGCAGTTGACCTGTGTGCAGC-3’; Reverse: 5’-CCCTCTAGGTCCAGTGGGTAAC-3’. PCR products were sequenced using the sanger method with a forward primer hybridizing in exon 5: 5’-CCACTGCCAAGGAGATCA-3’ (Figure S1A). Sequences are available upon request. Briefly, this variant causes a skipping of the entire exon 6 in the mRNA leading to a frameshift and a premature stop codon, thus undergoing nonsense-mediated mRNA decay as shown in Figure S1C. Consequently, a strong decrease of the corresponding mRNA steady state level is observed (Figure S1B). This MM variant was deposited in the *ClinVar* database ([VCV000981372.1](https://www.ncbi.nlm.nih.gov/clinvar/variation/981372/)).

### RiboMethSeq

RiboMethSeq analysis of human LCL tRNAs was performed as described in (Marchand *et al*, 2017). Briefly, tRNAs extracted from LCLs were fragmented in 50 mM bicarbonate buffer pH 9.2 for 15 minutes at 95°C. The reaction was stopped by ethanol precipitation. The pellet was washed with 80% ethanol and sizes of generated RNA fragments were assessed by capillary electrophoresis using a small RNA chip on Bioanalyzer 2100 (Agilent, USA). RNA fragments were directly 3’-end dephosphorylated using 5 U of Antarctic Phosphatase (New England Biolabs, UK) for 30 minutes at 37°C. After inactivation of the phosphatase for 5 minutes at 70°C, RNA fragments were phosphorylated at the 5’-end using T4 PNK and 1 mM ATP for one hour at 37°C. End-repaired RNA fragments were then purified using RNeasy MinElute Cleanup kit (QIAGEN, Germany) according to the manufacturer’s recommendations. RNA fragments were converted to library using NEBNext® Small RNA Library kit (ref#E7330S, New England Biolabs, UK) following the manufacturer’s instructions. DNA library quality was assessed using a High Sensitivity DNA chip on a Bioanalyzer 2100. Library sequencing was performed on Illumina HiSeq 1000 in single-read mode for 50 nt. Primary analysis of sequencing quality was performed with RTA 2.12 software, to insure > Q30 quality score for > 95 % of obtained sequences.

Following SR50 sequencing run, demultiplexing was performed with BclToFastq v2.4, reads not passing quality filter were removed. Raw reads after demultiplexing were trimmed with Trimmomatic v0.32 (Bolger *et al*, 2014). Alignment to the reference tDNA sequences was performed with bowtie 2 ver2.2.4 (Langmead *et al*, 2009) in End-to-End mode. Uniquely mapped reads were extracted from *.sam file by RNA ID and converted to *.bed format using bedtools v2.25.0 (Quinlan, 2014). Positional counting of 5’-and 3’-ends of each read was performed with awk Unix command. Further treatment steps were performed in R environment (v3.0.1). In brief, 5’-and 3’-end counts were merged together by RNA position and used for calculation of ScoreMEAN (derived from MAX Score (Pichot *et al*, 2020), as well as Scores A and B (Birkedal *et al*, 2015) and MethScore (Score C) (Marchand *et al*, 2016). Scores were calculated in the window of −2 to +2 neighbouring nucleotides. Profiles of RNA cleavage at selected (candidate and previously known) positions were extracted and visually inspected.

Analysis of human tRNA 2’-O-methylation by RiboMethSeq was performed using the optimised non-redundant collection of reference tRNA sequences. This reduced collection contains 43 tRNA species and was validated by analysis of several experimentally obtained RiboMethSeq sequencing datasets (Pichot *et al*, 2021). Alignment of RiboMethSeq reads obtained in this study also confirmed low content in ambiguously mapped reads. In order to establish a reliable map of Nm positions in human tRNA anticodon loop, RiboMethSeq cleavage profiles were used to calculate detection scores (Mean and ScoreA2) (Pichot *et al*, 2020). However, this scoring strategy shows its limits in the case of short and highly structured RNAs (like tRNAs), since the cleavage profile is highly irregular. In addition, since these scores are calculated for 2 neighbouring nucleotides, simultaneous loss of two closely located Nm residues (*e.g*. Cm_32_ and Gm_34_ in tRNA^Phe^) makes analysis of raw score misleading (Angelova *et al*, 2020). Moreover, the presence of multiple RT-arresting modifications (Anreiter *et al*, 2021) in the same tRNAs (m^1^A, m^1^G, m^2^2G, m^3^C, etc) reduces coverage in the upstream regions. Considering all these limitations, visual inspection of raw cleavage profiles revealed to be the most appropriate, since changes in protection of a given nucleotide represent modulation of its Nm methylation status. Analysis of alignment statistics demonstrated that the majority of human tRNAs are well represented in the analysed datasets and proportion of uniquely mapped reads were >90% for all tRNA sequences, except tRNALeu(CCA) family, composed of 3 highly similar species. Only limited coverage of totally mapped reads <7500 reads/tRNA (∼100 reads/position) was obtained for 5 tRNAs (Arg_TCG, Leu_CAA2, Ser_CGA_TGA1, Thr_CGT and Tyr_ATA).

In order to identify potential Nm32/Nm34 residues, raw cleavage profiles of the 11 nt region around pos 33 were visually inspected and profiles for WT samples were compared to *FTSJ1* mutants. Due to the limited number of mapped raw reads, coverage in the anticodon loop for Leu_CAA, Ser_CGA_TGA1, Thr_CGT and Tyr_ATA was insufficient; thus, these species were excluded from further analysis. The results of this analysis are given in Table 2. This analysis allowed to identify 10 Nm32 and 4 Nm34 modifications on tRNAs ACL. Inosine residues formed by deamination of A34 at the wobble tRNA position (FTSJ1-independent) are visible in the sequencing data and are also shown in Table 2. 10 Nm32 and 3 Nm34 modifications were found to be FTSJ1-dependent. The only exception is Cm34 in tRNAMet_CAT known to be formed by snoRNA-guided Fibrillarin (Vitali & Kiss, 2019). Comparison of these data with previously reported Nm modifications in human tRNA anticodon loop demonstrated that 2/3 of the observed sites have been described, either in tRNAdb2009 ((Jühling *et al*, 2009), http://trnadb.bioinf.uni-leipzig.de/), or in two recent studies used LC-MS/MS analysis (Nagayoshi *et al*, 2021; Li *et al*, 2020b). Table 2 also shows those modifications in other organisms including yeast, mice and *Drosophila*. We were not able to confirm Nm residues previously reported in tRNASec_TCA (Nm34) and tRNAVal_AAC(Cm32), however, due to sequence similarity, tRNAVal_AAC clusters together with two other tRNAVal (CAC and TAC1). tRNALeu_AAG and Leu_TAG have similar sequences and thus were not distinguished by sequencing, however Nm32 was detected.

**Table 1.** FTSJ1 targets tRNA^Phe^ at positions 32 and 34 in humans. Control and affected FTSJ1 individuals Nm status at positions 32 and 34 of human tRNA^Phe^.

| tRNA<sup>Phe(GAA)</sup> |  |  |  |
| --- | --- | --- | --- |
| Individual | Cm<sub>32</sub> | Gm<sub>34</sub> | LCL code name |
| Control individuals | Present | Present | LCL16<br>LCL18<br>LCL24<br>LCL54 |
| Affected individuals with FTSJ1 variant | Absent | Absent | LCL65AW<br>LCL65JW<br>LCL11<br>LCLMM |
| Affected individual with FTSJ1 variant | Present | Absent | LCL22 |

**Table 2.** FTSJ1 targets multiple human tRNAs at positions 32 and 34. A summary of tRNA nucleotides revealed to date, including by the current study, as targets of human FTSJ1, as well as those targeted by *Drosophila* Trm7_32 and Trm7_34, and yeast Trm7 in the respective organisms. For the tRNA targets are given the isotype (determined by the bound amino acid) and the isoacceptor (determined by the ACL sequence). In blue are highlighted the studies done with the site-specific RiboMethSeq and in grey, the ones done by mass spectrometry (MS) single nucleotide analysis. n.d. stands for non-determined and indicates that the tRNA was not tested or if tested the data was not analysable. no stands for non-detected Nm. Cm, Gm and Um stand for 2’-O-methylated respectively C, G and U nucleotides. * indicates Nm RiboMethSeq detection by visual inspection of the raw reads profile not MethScore, see Figure S1D for an example. When several anticodon sequences are present for tRNA isoacceptors, proportion of every sequence in the healthy subject is indicated on the bottom. Cm^#^ indicates Cm detection in (Li *et al*, 2020b) that could be due to a high sequence similarity with another tRNAArg, tRNAArg(CCG)-2-1 containing a C32. The observed Cm decrease in FTSJ1 KO cells in this study may come from C32 of tRNAArg(CCG)-2-1 that was modified by FTSJ1 and not from the C34 level of tRNAArg(CCG). Cm^@^ indicates hm5Cm34 or f5Cm34 in tRNALeu(CAA) shown in (Kawarada *et al*, 2017). **I** stand for inosine (FTSJ1 independent). U^?^m* indicates clear FTSJ1-dependence, however, the exact nature of this modified U remains unknown. tRNAArg (UCG) and (CCG) have identical sequences but differ only at positions 32 and 34. $ stand for UCG isodecoders (sequences in the Material and Methods section). tRNALeu (A/IAG) and (UAG) are isoacceptors, they differs only by the N34 nucleotide, and both have Um32 (or ψm32).

| tRNA target | *Human* |  |  |  | *Drosophila* |  | *S. cerevisiae* |  | *Mouse* |  |
| --- | --- | --- | --- | --- | --- | --- | --- | --- | --- | --- |
|  | Current RiboMethSeq |  | Previous HPLC/ MS |  | Previous RiboMethSeq |  | Previous HPLC and/or MS |  | Previous HPLC/ MS |  |
|  | *N32* | *N34* | *N32* | *N34* | *N32* | *N34* | *N32* | *N34* | *N32* | *N34* |
| Arg (UCG1)\$ | **Cm** | no | **Cm**<br>(Li *et al*, 2020b) | **no** | no | no | n.d. | n.d. | n.d. | n.d. |
| Arg (CCG) | **Um** | no | **Um, Cm**<br>(Li *et al*, 2020b) | **Cm<sup>#</sup>**<br>(Li *et al*, 2020b) | no | no | n.d. | n.d. | n.d. | n.d. |
| Arg (ACG) | **Cm\*** | **I** | **Cm**<br>(Li *et al*, 2020b) | no | **Cm**<br>(Angelova & Dimitrova *et al*, 2020) | no | n.d. | n.d. | **Cm**<br>(Nagayoshi *et al*, 2021) | n.d. |
| Arg (UCG2)\$ | **Cm** | no | n.d. | n.d. | n.d. | n.d. | n.d. | n.d. | n.d. | n.d. |
| Leu (CAG_CAA)<br>91%_9% | **Um** | **no** | no | **Cm<sup>@</sup>**<br>(Li *et al*, 2020b; Kawarada *et al*, 2017) | **Cm**<br>(Angelova & Dimitrova *et al*, 2020) | **Cm**<br>(Angelova & Dimitrova *et al*, 2020) | n.d. | n.d. | n.d. | **Um_hm<sup>5</sup>Cm**<br>(Nagayoshi *et al*, 2021) |
| Leu (UAA) | no | **U<sup>2</sup>m\*** | n.d. | no | no | no | **Cm**<br>(Guy *et al*, 2012b) | ncm<sup>5</sup>**Um**<br>(Glasser *et al*, 1992; Guy *et al*, 2012b) | **Cm**<br>(Nagayoshi *et al*, 2021) | ncm<sup>5</sup>**Um**<br>(Nagayoshi *et al*, 2021) |
| Leu (A/IAG)<br>76% | **U/ψm** | **I** | n.d. | no | no | no | n.d. | n.d. | **ψm**<br>(Nagayoshi *et al*, 2021) | n.d. |
| Leu (UAG)<br>24% | **U/ψm** | **no** | n.d. | no | no | no | n.d. | n.d. | **Um**<br>(Nagayoshi *et al*, 2021) | n.d. |
| Phe (GAA) | **Cm\*** | **Gm** | **Cm**<br>(Guy *et al*, 2015; Li *et al*, 2020b; Nagayoshi *et al*, 2021) | **Gm**<br>(Guy *et al*, 2015; Li *et al*, 2020b; Nagayoshi *et al*, 2021) | **Cm\***<br>(Angelova & Dimitrova *et al*, 2020) | **Gm**<br>(Angelova & Dimitrova *et al*, 2020) | **Cm**<br>(Guy *et al*, 2012b) | **Gm**<br>(Guy *et al*, 2012b) | **Cm**<br>(Nagayoshi *et al*, 2021) | **Gm**<br>(Nagayoshi *et al*, 2021) |
| Trp<br>(CCA) | Cm | Cm | Cm<br>(Guy *et al*, 2015; Li *et al*, 2020b; Nagayoshi *et al*, 2021) | Cm<br>(Guy *et al*, 2015; Li *et al*, 2020b; Nagayoshi *et al*, 2021) | Cm<br>(Angelova & Dimitrova *et al*, 2020) | Cm<br>(Angelova & Dimitrova *et al*, 2020) | Cm<br>(Guy *et al*, 2012b) | Cm<br>(Guy *et al*, 2012b) | Cm<br>(Nagayoshi *et al*, 2021) | Cm<br>(Nagayoshi *et al*, 2021) |
| Gln<br>(CUG_UUG)<br>92%_8% | Cm | no | Cm<br>(Li *et al*, 2020b) | n.d. | Cm<br>(Angelova & Dimitrova *et al*, 2020) | no | n.d. | n.d. | Cm<br>(Nagayoshi *et al*, 2021) |  |
| Gly<br>(CCC) | Um | no | n.d. | no | no | no | n.d. | n.d. | n.d. | n.d. |
| Val<br>(AAC_CAC_TAC)<br>73%_26%_1% | no | I<br>(AAC) | Cm<br>(Nagayoshi *et al*, 2021) | n.d. | Cm<br>(Angelova & Dimitrova *et al*, 2020) | no | n.d. | n.d. | Cm<br>(Nagayoshi *et al*, 2021) |  |
| Pro<br>(AGG_CGG_UGG)<br>34%_23%_42% | Um* | I<br>(AGC) | no | n.d. | n.d. | n.d. | n.d. | n.d. | n.d. | n.d. |
| Cys<br>(GCA_ACA)<br>97%_3% | Cm* | no | n.d. | n.d. | n.d. | n.d. | n.d. | n.d. | n.d. | n.d. |
| Met<br>(CAU)<br>*non FTSJ1 Target* | no | Cm | no | Cm<br>(Vitali & Kiss, 2019; Li *et al*, 2020b) | no | no | no | no | n.d. | n.d. |

### mRNA sequencing and data analysis

mRNA sequencing was performed as in (Khalil *et al*, 2018). 5 μg of total RNA were treated by 1MBU of DNAse (BaseLine-Zero™ DNAse, Epicentre, USA) for 20 min at 37°C to remove residual genomic DNA contamination. RNA quality was verified by PicoRNA chip on Bioanalyzer 2100 (Agilent, USA) to ensure RIN (RNA Integrity Number) > 8.0. PolyA + fraction was isolated from 4.5 μg of DNAse-treated total RNA using NEBNext Oligo d(T)25 Magnetic beads kit (NEB, USA), according to manufacturer’s recommendations. PolyA + enrichment and the absence of residual rRNA contamination were verified using PicoRNA chips on Bioanalyzer 2100 (Agilent, USA). PolyA + fraction (1 ng for each sample) was used for whole-transcriptome library preparation using ScriptSeq v2 RNA-Seq kit (Illumina, USA). Libraries amplified in 14 PCR cycles were purified using Agencourt AMPure XP beads (Beckman-Coulter, USA), at a ratio 0.9x to remove adapter dimer contamination. Quality of the libraries was verified by HS DNA Chip on Bioanalyzer 2100 (Agilent, USA) and quantification done by Qubit 2.0 with appropriate RNA quantification kit. Sequencing was performed on HiSeq1000 (Illumina, USA) in single read SR50 mode. About 50 million of raw sequencing reads were obtained for each sample. Adapters were trimmed by Trimmomatic v0.32 (Bolger *et al*, 2014) and the resulting sequencing reads aligned in sensitive-local mode by Bowtie 2 v2.2.4 (Langmead & Salzberg, 2012) to hg19 build of human genome. Differential expression was analyzed using *.bam files in DESeq2 package (Love *et al*, 2014) under R environment. Analysis of KEGG and Gene Ontology pathways for differentially expressed genes was done under R environment.

### small RNA sequencing and data analysis

Small RNA-Seq libraries were generated from 1000 ng of total RNA using TruSeq Small RNA Library Prep Kit (Illumina, San Diego, CA), according to manufacturer’s instructions. Briefly, in the first step, RNA adapters were sequentially ligated to each end of the RNA, first the 3′ RNA adapter that is specifically modified to target microRNAs and other small RNAs, then the 5’ RNA adapter. Small RNA ligated with 3′ and 5′ adapters were reverse transcribed and PCR amplified (30 sec at 98°C; [10 sec at 98°C, 30 sec at 60°C, 15 sec at 72°C] × 13 cycles; 10 min at 72°C) to create cDNA constructs. Amplified cDNA constructs of 20 to 40 nt were selectively isolated by acrylamide gel purification followed by ethanol precipitation. The final cDNA libraries were checked for quality and quantified using capillary electrophoresis and sequenced on the Illumina HiSeq 4000 at the Institut de Génétique et de Biologie Moléculaire et Cellulaire (IGBMC) GenomEast sequencing platform.

For small RNA data analysis, adapters were trimmed from total reads using FASTX_Toolkit [http://hannonlab.cshl.edu/fastx_toolkit/]. Only trimmed reads with a length between 15 and 40 nucleotides were kept for further analysis. Data analysis was performed according to published pipeline ncPRO-seq (Chen *et al*, 2012). Briefly, reads were mapped onto the human genome assembly hg19 with Bowtie v1.0.0. The annotations for miRNAs were done with miRBase v21. The normalization and comparisons of interest were performed using the test for differential expression, proposed by (Love *et al*, 2014) and implemented in the Bioconductor package DESeq2 v1.22.2 [http://bioconductor.org/]. MicroRNA target prediction was performed using miRNet 2.0 (Chang *et al*, 2020).

### Northern blotting

For northern blotting analysis of tRNA, 5 μg of total RNA from human LCLs were resolved on 15 % urea-polyacrylamide gels for approximately 2h in 0.5x TBE buffer at 150 V, then transferred to Hybond-NX membrane (GE Healthcare) in 0,5× TBE buffer for 1h at 350 mA of current and EDC-cross-linked for 45 min at 60°C with a solution containing 33 mg/ml of 1-ethyl-3-(3-dimethylaminopropyl)carbodiimide (EDC) (Sigma Aldrich), 11 ng/ul of 1-methylimidazol and 0.46% of HCl. The membranes were first pre-hybridized for 1h at 42°C in a hybridization buffer containing 5xSSC, 7% SDS, 5.6 mM NaH_2_PO_4_, 14.4 mM Na_2_HPO_4_ and 1× Denhardt’s solution. DNA oligonucleotide probes were labelled with ^32^P at the 5’-end by T4 polynucleotide kinase following manufacturer’s instructions (Fermentas). The membranes were hybridized with the labelled probes overnight at 42°C in the hybridization buffer, then washed twice for 15 min in wash buffer A (3x SSC and 5% SDS) and twice in wash buffer B (1× SSC and 1% SDS) before film exposure at −80°C for variable time durations. Probe sequences are available in the *Primers and Probes section*.

### RT-qPCR

RNA was extracted from human LCLs using TRI-Reagent (Sigma Aldrich). After DNase digestion of total RNA using the TURBO DNA-free™ Kit (Ambion), 1 μg was used in a reverse transcription reaction with Random Primers (Promega) and RevertAid Reverse Transcriptase (ref. EP0442, Thermofisher). The cDNA was used to perform qPCR on a CFX96 Touch™ Real-Time PCR Detection System (Bio Rad) using target-specific primers. *hGAPDH* was used for normalization (*Primers and Probes section*). The analysis was performed using ΔΔ Ct, on three biological replicates. Statistical analysis using a bilateral Student’s t-test was performed and *p*-values were calculated.

### NMD inhibition test

LCLs were seeded in 25 cm cell culture plates at a density of 3.10^6^ cells and treated with 100 μg/mL of cycloheximide or equal volume of water as a control for six hours. Cells were harvested by centrifugation at 1000 rpm for 5min and flash frozen in liquid nitrogen. RNA extraction was carried out using TRI-reagent (Sigma Aldrich) following the supplier’s protocol. DNAse I digestion was carried out using RNAse free DNase I (M0303S- NEB), and reverse transcription on 1 μg of DNase treated total RNA was performed using RevertAid Reverse Transcriptase. Quantitative PCR was performed as specified above using specific primers for *FTSJ1* and *GAPDH*.

### miRNA complementation experiments

mirVana™ miRNA Mimics and Inhibitors were used for hsa-miR-181a-5p overexpression/inhibition (Ambion™ - 4464066 and 4464084). HeLa cells were transfected with corresponding mirVana™ miRNA in 24 well plates at a density of 20.000 cells per well, using Lipofectamine™ RNAiMAX (CAT# 13778100-Invitrogen™). We set up the transfection ratios to 15 pmol of miRNA mimic/μL of Lipofectamine™, and 30 pmol of miRNA inhibitor/μL of Lipofectamine™. Cells were harvested 48 hours post-transfection and assayed for target gene expression. miRNA quantification was performed by RTqPCR on miR181a-5p using Qiagen’s miRCURY LNA miRNA PCR System. Reverse transcription is performed using miRCURY LNA RT Kit (339340) and qPCR using miRCURY LNA SYBR® Green PCR Kit (339346). LNA enhanced primers were used for miRNA Sybr green qPCR (Refer to the list of primers and probes).

### Primers, Probes and Sequences

Northern blot analysis was performed using *hsa-miR-181a-5p* specific probes with the following sequences: 5’-AACATTCAACGCTGTCGGTGAGT-3’ (sense probe) and 5’-ACTCACCGACAGCGTTGAATGTT-3’ (antisense probe). Human U6 specific probe was used for detecting U6 as a loading control: 5’-GCAAGGATGACACGCAAATTCGTGA-3’ (sense probe) and 5’-TCACGAATTTGCGTGTCATCCTTGC-3’ (antisense probe). qPCR analysis (after an RT reaction performed with random primers) were performed with the use of primers with the following sequences:

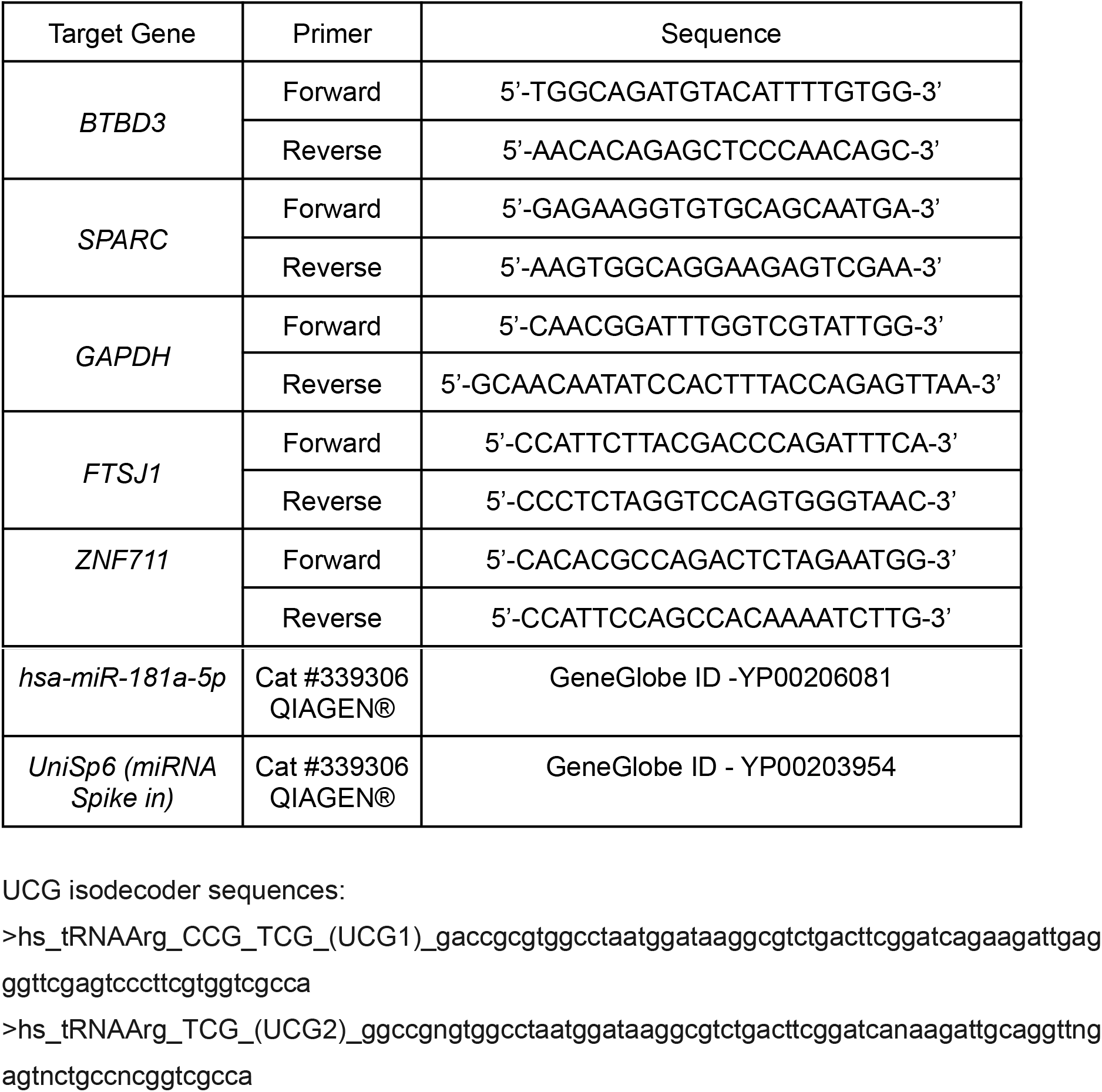

UCG isodecoder sequences:

>hs_tRNAArg_CCG_TCG_(UCG1)_gaccgcgtggcctaatggataaggcgtctgacttcggatcagaagattgagggttcgagtcccttcgtggtcgcca

>hs_tRNAArg_TCG_(UCG2)_ggccgngtggcctaatggataaggcgtctgacttcggatcanaagattgcaggttngagtnctgccncggtcgcca

### iPSC culture and maintenance

iPSCs cell line WTSli002 purchased from EBISC (European bank for induced pluripotent cells) were maintained on feeder-free conditions on Geltrex LDEV-Free hESC-qualified Reduced Growth Factor Basement Membrane Matrix (ThermoFisher Scientific, A1413302) in Essential 8™ Flex Media Kit (ThermoFisher Scientific, A2858501) with 0,1% Penicillin/Streptomycin (ThermoFisher Scientific, 15140122).

### iPSC differentiation in dorsal NPCs

To obtain Neural progenitor cells (NPCs) from the dorsal telencephalon, embryoid bodies (EB) were formed by incubating iPSCs clusters with Accutase (ThermoFisher Scientific, A1110501) for 7 min at 37°C and dissociated into single cells. To obtain EB of the same size, 3 × 10^6^ cells were added per well in the AggreWell 800 plate (STEMCELL Technologies, 34815) with Essential 8™ Flex Media supplemented with Stemgent hES Cell Cloning & Recovery Supplement (1X, Ozyme, STE01-0014-500) and incubated at 37°C with 5% CO_2_ (Day-1). After 24 hours in culture (Day0), EB from each microwell were collected by pipetting up and down the medium several times and transferred into Corning® non-treated culture dishes (Merck, CLS430591-500EA) in EB medium containing DMEM/F12 GlutaMAX (ThermoFisher Scientific, 35050061), 20% KnockOut™ Serum Replacement (ThermoFisher Scientific, 10828028), 1% Non-Essential Amino Acid (ThermoFisher Scientific,11140035), 0,1% Penicillin/Streptomycin (ThermoFisher Scientific, 15140122), 100 μM 2-mercaptoethanol (ThermoFisher Scientific, 31350010), supplemented with two inhibitors of the SMAD signalling pathway, 2,5 μM Dorsomorphin (Sigma-Aldrich, P5499) and 10 μM SB-431542 (Abcam, ab120163). EB medium supplemented as described previously was changed every day for 5 days. On Day 6, floating EBs are plated on 0,01 % Poly-L-ornithine (Sigma-Aldrich, P4957) and 5 μg/mL Laminin (Sigma-Aldrich, L2020) coated dishes for rosette expansion in Neurobasal minus vitamin A (ThermoFisher Scientific 10888), B-27 supplement without vitamin A (ThermoFisher Scientific 12587), 1% GlutaMAX (ThermoFisher Scientific 35050061), 0,1% Penicillin/Streptomycin (ThermoFisher Scientific 15140122) and 100 μM 2-Mercaptoethanol (ThermoFisher Scientific 31350010). The neural medium was supplemented with 10 ng/mL epidermal growth factor (PreproTech AF-100-15) and 10 ng/mL basic fibroblast growth factor (R&D Systems 234-FSE-025). From day 6 to day 10 the medium was changed everyday until the appearance of rosettes. On day 10, rosettes are manually picked up using a syringe and dissociated with Accutase, then seeded on Poly-L-ornithine/Laminin coated dishes for expansion of dorsal NPCs. They were maintained with passage for two additional weeks to achieve a large pool of neural precursor cells (NPCs).

### NPC drug treatment

NPCs are seeded in Poly-L-Ornithine and Laminin coated coverslips in 24 well plates at a density of 2.10^5^ cells per well. After 48 hours, the medium is changed and combined with 100 μM of 2,6 Diaminopurine (DAP) (Sigma Aldrich 247847) or equal volume of sterile H_2_O.

### NPC immunostainings

24 hours after DAP treatment NPCs were fixed in 4% paraformaldehyde for 10 min, permeabilized and blocked for 45 minutes with blocking buffer (PBS supplemented with 0.3% Triton-X100, 2% horse serum). Primary antibodies, Sox2 (1/500, Milipore AB5603) and DCX (1/2000, Milipore AB2253), were incubated overnight at 4°C using the same solution. Cells were rinsed three times with PBS and incubated 1 hour at RT with secondary antibodies and DAPI (1/10000, Sigma-Aldrich D9564) diluted in the same solution and rinsed 3 times with PBS before mounting on slides with VectaShield® Vibrance mounting medium.

### Neuronal cells image acquisitions

Images were acquired in z-stacks using a confocal microscope Nikon A1R HD25 with a 60X objective. Images were flattened with a max intensity Z-projection.

### Neurogenesis quantification

All cells (DAPI) from each acquisition were numbered using Fiji’s point tool. Cells expressing DCX (immature neurons) and SOX2 (NPCs and intermediates which also started expressing DCX) were also numbered on 5 to 6 microscopy images. Over 1400 cells were numbered for each condition in triplicate. A ratio of DCX expressing cells is calculated over the total cell number and expressed in fold change and compared between DAP treated and untreated cells.

### Branching quantifications

All DCX expressing neurons were traced using Simple Neurite Tracer (SNT) from the Neuroanatomy Plugin by Fiji. Length measurements of traces were performed using the SNT Measure Menu, and thin projections were counted manually using Fiji’s point tool. Quantifications were performed on 5 acquisitions and each IF experiment was done in triplicate. Ratios for the number of thin projections/neuron length (mm) were calculated and compared between DAP treated and control cells.

### *Drosophila* NMJ analysis

For NMJ staining, third instar larvae were dissected in cold PBS and fixed with 4% paraformaldehyde in PBS for 45 min. Larvae were then washed in PBST (PBS + 0.5% Triton X100) six times for 30 min and incubated overnight at 4°C with mouse anti-synaptotagmin, 1:200 (3H2 2D7, Developmental Studies Hybridoma Bank, DSHB). After six 30 min washes with PBST, secondary antibody anti-mouse conjugated to Alexa-488 and TRITC-conjugated anti-HRP (Jackson ImmunoResearch) were used at a concentration of 1:1,000 and incubated at room temperature for 2 h. Larvae were washed again six times with PBST and finally mounted in Vectashield (Vector Laboratories).

For DAP treatment, freshly hatched *Canton-S* flies were collected and placed on a normal food medium containing 600 μM of 2,6 Diaminopurine (DAP) (Sigma aldrich 247847). After 5 days, third instar larvae were dissected and subjected to NMJ staining.

Images from muscles 6–7 (segment A2–A3) were acquired with a Zeiss LSM 710 confocal microscope. Serial optical sections at 1,024 × 1,024 pixels with 0.4 μm thickness were obtained with the ×40 objective. Bouton number was quantified using Imaris 9 software. ImageJ software was used to measure the muscle area and the NMJ axon length and branching. Statistical tests were performed in GraphPad (PRISM 8).

### *Drosophila* behaviour assays

Flies were raised at 25°C for associative memory assays and the corresponding controls. All behaviour experiments were performed on young adults (1-3 day-old). All behaviour experiments were performed on starved flies, which is a prerequisite for appetitive conditioning with a sucrose reinforcement. 0-2 days after hatching, flies were put on starvation for 21h at 25°C on mineral water (Evian). **Appetitive memory assay:** Appetitive associative conditioning was performed in custom-designed barrel-type apparatus as previously described (Colomb *et al*, 2009), which allows the parallel conditioning of three groups of flies. The odorants 3-octanol and 4-methylcyclohexanol, diluted in paraffin oil at a final concentration of 0,29 g·L^-1^, were used for conditioning and for the test of memory retrieval. Groups of 20–50 flies were subjected to one cycle of appetitive olfactory conditioning as follows: throughout the conditioning protocol, flies were submitted to a constant air flow at 0,6 L·min^-1^. After 90 s of habituation, flies were first exposed to an odorant (the CS^+^) for 1 min while given access to dried sucrose; flies were then exposed 45 s later to a second odorant without shocks (the CS^-^) for 1 min. 3-octanol and 4-methylcyclohexanol were alternately used as CS^+^ and CS^-^. The memory test was performed in a T-maze apparatus. Each of the two arms of the T-maze were connected to a bottle containing one odorant (either 3-octanol or 4-methylcyclohexanol) diluted in paraffin oil. The global air flow from both arms of the T-maze was set to 0,8 L·min^-1^. Flies were given 1 min in complete darkness to freely move within the T-maze. Then flies from each arm were collected and counted. The repartition of flies was used to calculate a memory score as ^(N^CS+^-N^CS-^)/(N^CS+^+ N^CS-^)^. A single performance index value is the average of two scores obtained from two groups of genotypically identical flies conditioned in two reciprocal experiments, using either odorant as the CS^+^. Thus values of performance index range between −1 and +1, the value of 0 (equal repartition) corresponding to ‘no memory’. The indicated ‘n’ is the number of independent performance index values for each genotype. LTM performance was assessed 24 hrs (+/− 2 hrs) after conditioning, STM 1 hr (+/− 30 min) after conditioning. **Innate odor avoidance and sucrose attraction assay:** Innate sucrose preference was measured in a T-maze. Flies were given the choice for 1 min between one arm of the T-maze coated with dried sucrose, and one empty arm. There was no air flow in the T-maze for this assay. Flies were then collected from each arm and counted; an attraction index was calculated as (N_sucrose_-N_empty_)/(N_sucrose_+N_empty_). The side of the T-maze with sucrose was alternated between experimental replicates. Innate odor avoidance was measured in a T-maze. One arm of the T-maze was connected to a bottle containing the tested odorant (3-octanol or 4-methylcyclohexanol) diluted in paraffin oil, the other arm was connected to a bottle containing paraffin oil only. The global air flow from both arms of the T-maze was set to 0,8 L·min^-1^. Flies were given 1 min in complete darkness to freely move within the T-maze. Flies were then collected from each arm and counted; an avoidance index was calculated as (N_air_-N_odor_)/(N_air_+N_odor_). The side of the T-maze with odorant-interlaced air was alternated between experimental replicates. **Quantification and statistical analysis**: All data are presented as mean ± SEM. Performances from different groups (mutant and control) were statistically compared using one-way ANOVA followed by Tukey’s posthoc pairwise comparison between the mutant genotypes and the control group.

## RESULTS

### Comprehensive identification of human FTSJ1 tRNA targets

To identify new tRNA targets of human FTSJ1, we compared the Nm modification profiles of positions 32 and 34 for all detectable tRNA species in human LCLs obtained from control individuals (n=4) *vs*. LCLs obtained from individuals with ID harbouring loss-of-function and pathogenic variants in *FTSJ1* (n=5, from four unrelated families) (Table 1). Four of these affected individuals were already described and harbour distinct molecular defects: a splice variant leading to a premature stop codon (Freude *et al*, 2004) (LCL65AW and LCL65JW), a deletion encompassing *FTSJ1* and its flanking gene *SLC38A5* (Froyen *et al*, 2007) (LCL11), and a missense variant (p.Ala26Pro) affecting an amino acid located close to FTSJ1 catalytic pocket, resulting in the loss of Gm_34_, but not of Cm_32_ in human tRNA^Phe^ (Guy *et al*, 2015) (LCL22). The last individual was not reported nor characterised before. This patient presents mild ID and behavioural manifestations and harbours a *de novo* pathogenic variant affecting the consensus acceptor splice site of exon 6 (NM_012280.3: c.362-2A>T) (LCL-MM). This mutation leads to the skipping of exon 6 in the mRNA (r.362_414del) leading to a frameshift and a premature stop codon (p.Val121Glyfs*51) (Figure S1A). *FTSJ1* mRNA steady state level in LCL-MM was significantly reduced when compared to LCL from control individuals (Figure S1B). In addition, treating the LCL-MM cells with cycloheximide to block translation, and thus the nonsense mediated mRNA decay (NMD) pathway (Tarpey *et al*, 2007), led to an increase of *FTSJ1* mRNA abundance (Figure S1C). This result suggests that *FTSJ1* mRNA from LCL-MM cells is likely degraded *via* the NMD pathway.

To obtain a comprehensive picture of the Nm-MTase specificity for FTSJ1 *in vivo*, we performed RiboMethSeq analysis on LCLs isolated from affected individuals described above and compared with LCL from healthy individuals. RiboMethSeq allows tRNA-wide Nm detection based on random RNA fragmentation by alkaline hydrolysis followed by library preparation and sequencing ((Marchand *et al*, 2017) and Material and Methods). Using this approach, we could confirm the known FTSJ1 targets (*e.g*. tRNA^Phe(GAA)^ and tRNA^Trp(CCA)^) and assign the FTSJ1-deposited Nm modifications to their predicted positions in the ACL (C_32_ and N_34_, Figure 1). However, using only the MethScore calculation we could not detect a variation for Cm_32_ in tRNA^Phe(GAA)^. This scoring strategy shows its limits in some particular situation as MethScore is calculated for 2 neighbouring nucleotides, thus simultaneous loss of two closely located Nm residues (*e.g*. Cm_32_ and Gm_34_ in tRNA^Phe^) makes analysis of MethScore misleading (Angelova *et al*, 2020). Moreover, the presence of multiple reverse transcription (RT) arresting hyper-modification (*e.g*. m^1^G37/o2yW37 (Anreiter *et al*, 2021)) in the same tRNA regions impairs RT, thereby reducing the number of cDNAs spanning the ACL. Nevertheless, considering all these potential limitations when using only MethScore calculation, a visual inspection of raw cleavage profiles was performed (Figure S1D and Table 2) and revealed to be the most appropriate. When visualising raw reads count profile, reads’ ends number at position 33 (Cm_32_) of tRNA^Phe(GAA)^ was increased in *FTSJ1* mutated cells (Figure S1D), indicating a loss of Cm_32_ of tRNA^Phe(GAA)^ in *FTSJ1* mutated LCLs. Thus, using both MethScore (Figure 1) and visual inspection on all RiboMethSeq human tRNA sequences (Figure S1D) we were able to confirm known FTSJ1 tRNA targets and, importantly, discover new FTSJ1-dependent Cm_32_/Um_32_ modification in tRNA^Gly^, tRNA^Leu^, tRNA^Pro^ and tRNA^Cys^ (see Table 2 for isoacceptors details). Unexpectedly, Um_34_ in tRNA^Leu(UAA)^ also demonstrated clear FTSJ1-dependence, however, the exact nature of this modified nucleotide remains unknown (Table 2). In contrast, the protection signal observed at position 32 in human tRNA^Ala(A/IGC)^ is not FTSJ1-dependent and most likely results from ψm_32_ (visible in HydraPsiSeq (Marchand *et al*, 2022) profiling (Y.M. personal communication)) and not Um_32_.

**Figure 1.**
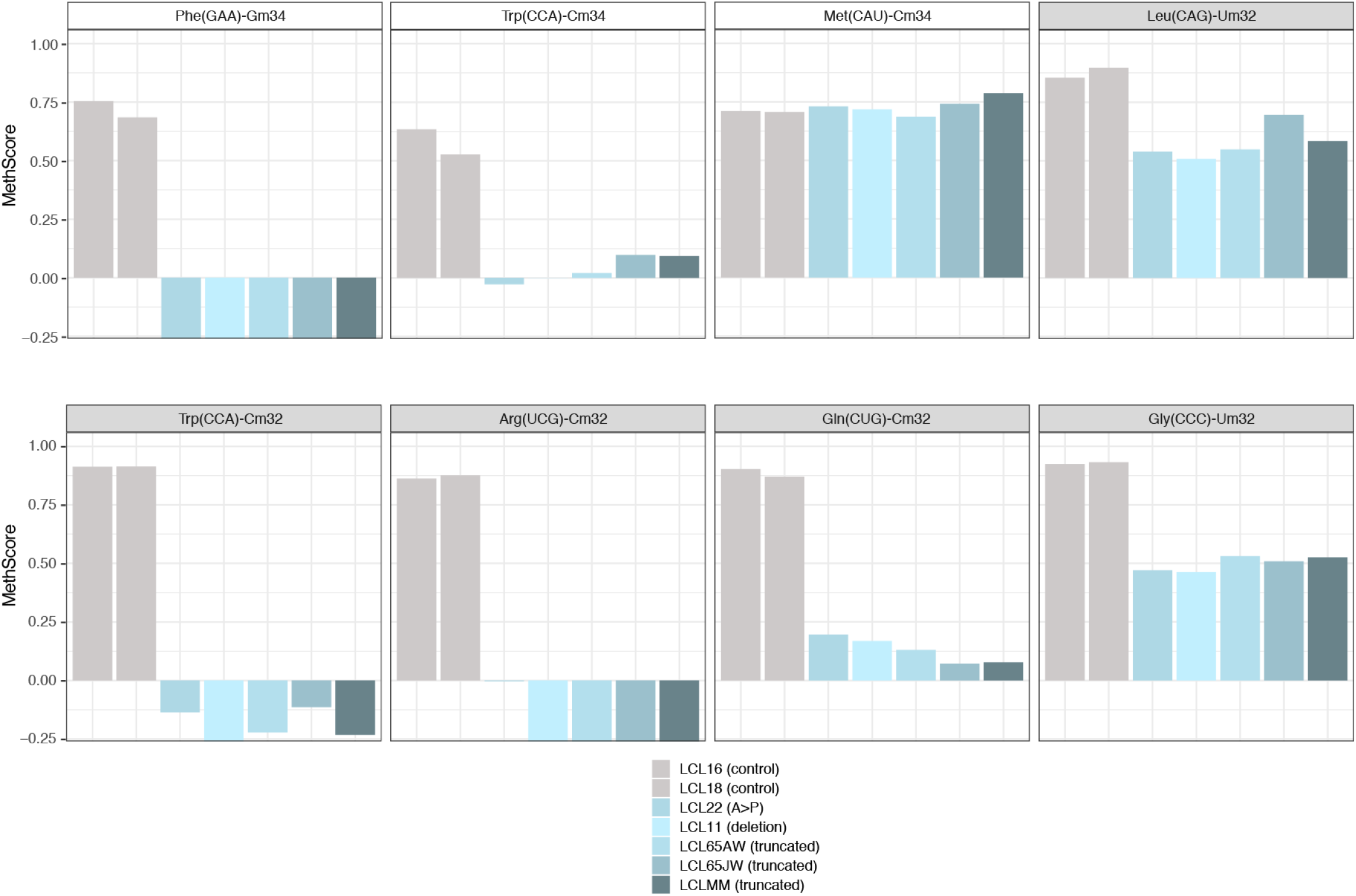
FTSJ1 targets multiple tRNAs at positions 32 and 34 in humans. Methylation scores (MethScore) for 2’-O-methylated positions in tRNAs showing altered methylation in FTSJ1 loss-of-function mutant LCLs. MethScore (Score C), representing the level of ribose methylation was calculated from protection profiles. Data are shown for positions 32 and 34 in different *H. sapiens* tRNAs as measured in different LCL lines that are indicated with different colour code. Grey: control LCL; blue: *FTSJ1* mutant LCLs. Met(CAU)-Cm34 is not deposited by FTSJ1 and shown here as a control (unaltered methylation in FTSJ1 mutants).

### *FTSJ1* loss of function deregulates mRNAs steady state level

To obtain insights into the impact of FTSJ1 loss on gene expression, we performed a transcriptome analysis in patient and control LCLs. Transcript differential expression analysis shows that FTSJ1 dysfunction led to a deregulation of 686 genes (Table 3 and Figures S2A and S2B). This relatively low number is in agreement with a previous report showing 775 genes deregulated in human HeLa cells knock-down for *FTSJ1* (Trzaska *et al*, 2020a), as well as with the 110 mRNAs deregulated in KD of one FTSJ1 *Drosophila* ortholog (Angelova and Dimitrova et al. 2020).

**Table 3.**
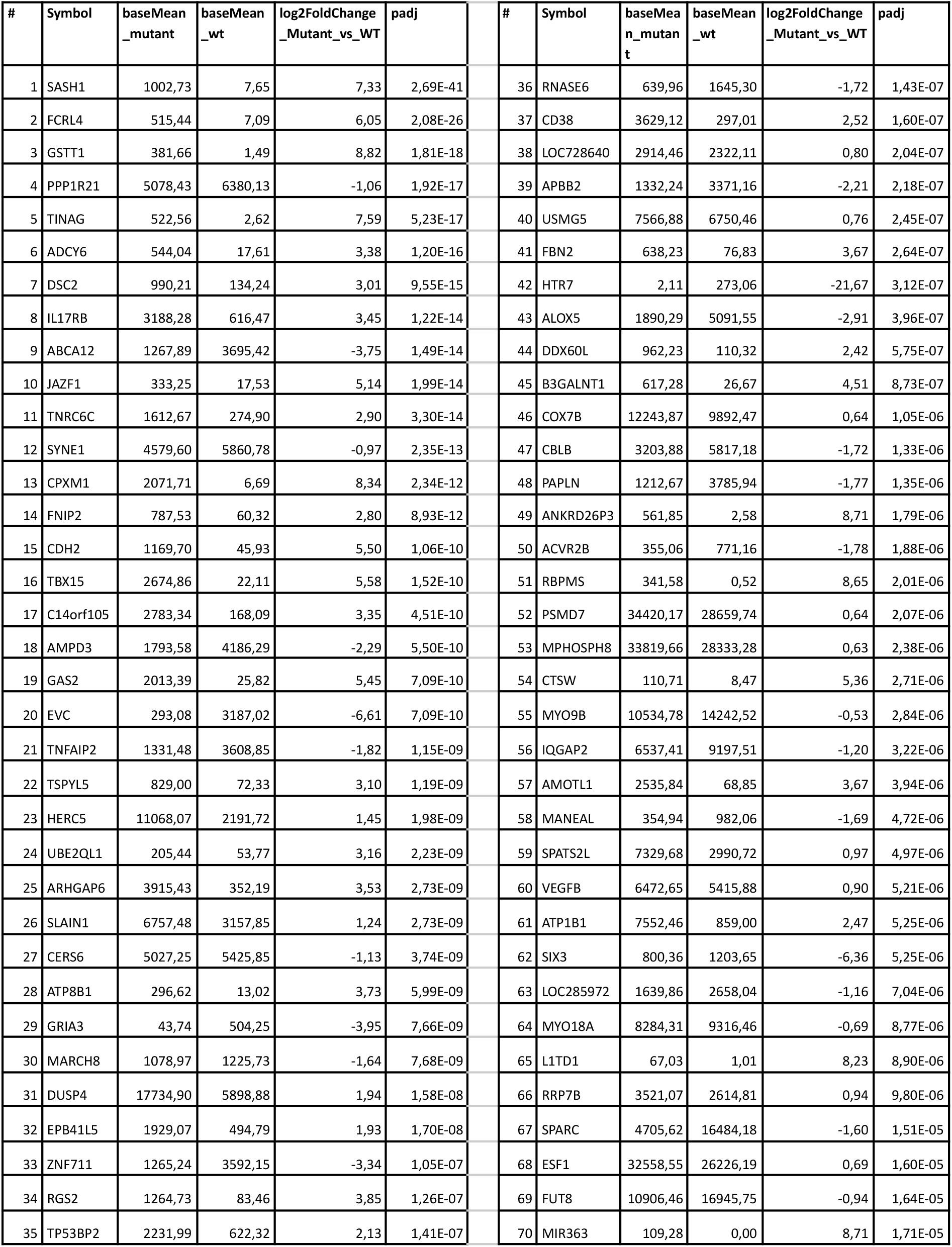
FTSJ1 loss of function leads to mRNAs deregulation in NSXLID affected individuals LCLs. A list of the 70 most significantly deregulated mRNAs in FTSJ1 LCLs mutants versus controls.

Even though LCLs do not have a neural origin, analysis of the genes deregulated in affected individuals revealed a clear enrichment (FE =7.9 with *p-*value =7.44E-06 and FDR =4.40E-03) in biological process Gene Ontology (GO) term corresponding to brain morphogenesis (Figure 2A). In addition, and similarly to what we reported in a previous mRNA-seq analysis of *Drosophila* S2 cells knocked-down for Trm7_34 (Angelova and Dimitrova et al. 2020), 5 out of the top 10 most enriched terms were related to mitochondrial biological processes. Also, in agreement with a recently described role of human FTSJ1 in translational control (Nagayoshi *et al*, 2021; Trzaska *et al*, 2020a) and of yeast Trm7 in the general amino-acid control pathway (Han *et al*, 2018), four biological processes related to translation were affected in *FTSJ1* mutated LCLs (FE >3.5, Figure 2A).

**Figure 2.**
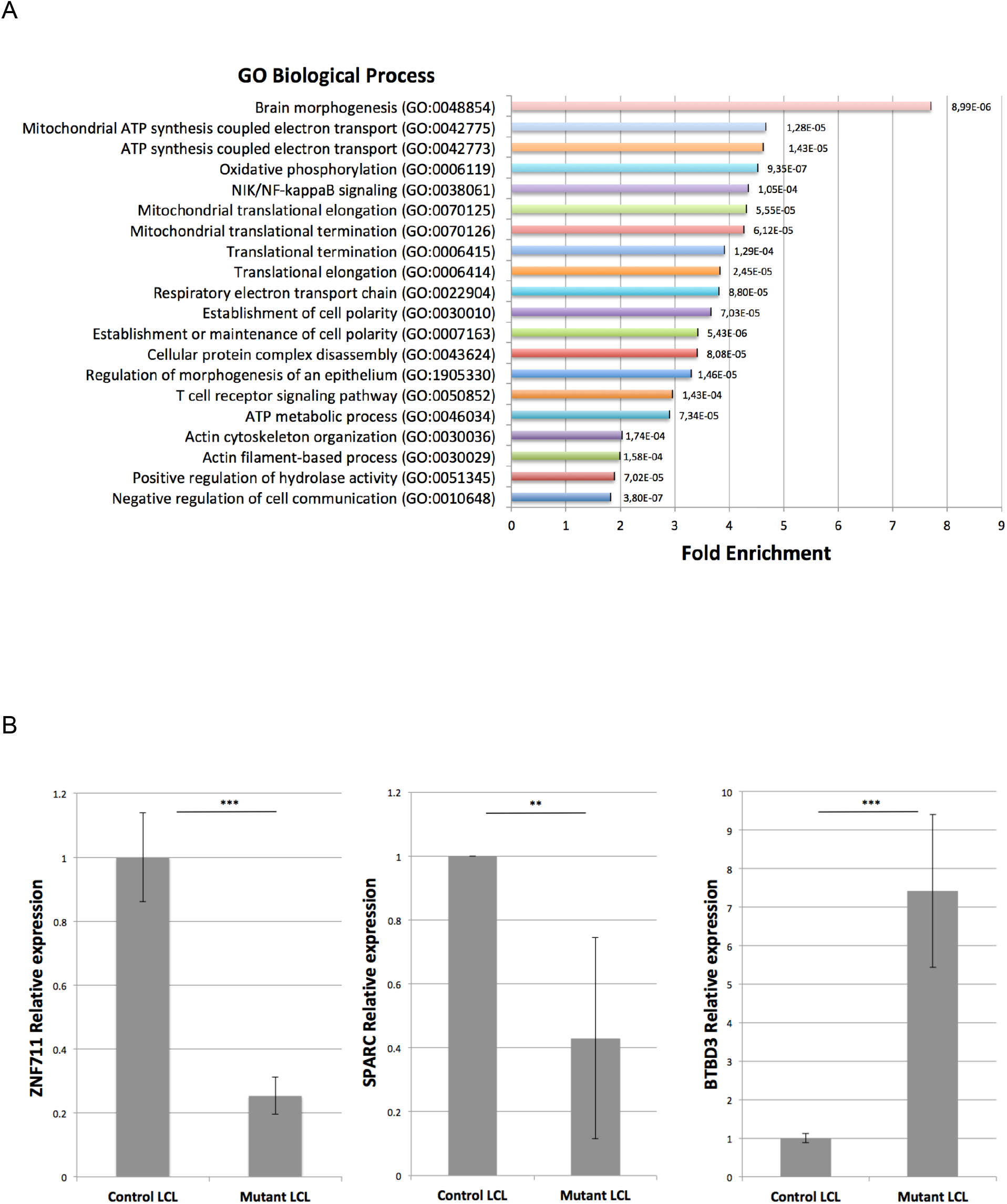
FTSJ1 loss of function leads to mRNAs deregulation in NSXLID affected individuals LCLs. **(A)** FTSJ1 loss of function mRNAs GO term. GO analysis of the 686 deregulated genes in FTSJ1 function-deficient LCLs derived from NSXLID affected individuals (5 mutants *vs*. 2 control LCLs; *p*-values are indicated with error bars on the right of each box. The most enriched GO term is brain morphogenesis. GO analysis was performed using http://geneontology.org/. **(B)** RT-qPCR analysis confirms deregulation in *ZNF711, BTBD3* and *SPARC* mRNAs expression levels. Normalized to *GAPDH* steady state levels. n>3. *p*-values were calculated with paired Student’s *t*-test \*\**p*<0,01, ***p<0,001. WT values: mean of 2 control FTSJ1 LCL. Mutant values: mean of all (×5) *FTSJ1* mutant LCLs of this study, or two (LCL MM and LCL 65JW) for *ZNF711* qPCR.

To strengthen the transcriptome analysis, we selected three representative and disease-relevant deregulated mRNAs based on their fold change level of expression and related involvement in brain or cancer diseases. Mutations in the human *ZNF711* gene were previously reported to be involved in the development of ID (van der Werf *et al*, 2017). The mRNA-seq and RT-qPCR analyses showed a significant downregulation of *ZNF711* mRNA in *FTSJ1* mutant LCLs when compared to control LCLs (Table 3 and Figure 2B). BTBD3 activity is known to direct the dendritic field orientation during development of the sensory neuron in mice cortex (Matsui *et al*, 2013) and to regulate mice behaviours (Thompson *et al*, 2019). We found that *BTBD3* mRNA was significantly upregulated in both mRNA-seq and RT-qPCR analyses (Figure 2B). Lastly, SPARC (Tai &Tang, 2008) and more recently FTSJ1 (Holzer *et al*, 2019; He *et al*, 2020) gene products activities were proposed to be involved in both metastasis and tumour suppression. In the absence of FTSJ1, we could confirm that *SPARC* mRNA was significantly reduced (Table 3 and Figure 2B). Taken together, these results show deregulation of some mRNAs linked to cancer and brain functioning in *FTSJ1* affected individuals’ blood derived LCLs.

### FTSJ1 loss of function affects the miRNA population

Our previous work on the *Drosophila* homologs of FTSJ1, Trm7_32 and Trm7_34, showed that their loss of functions led to perturbations in the small non-coding RNA (sncRNA) gene silencing pathways, including the miRNA population (Angelova and Dimitrova et al. 2020)To address whether such small RNA perturbations are conserved in NSXLID affected individuals we performed small RNA sequencing on the 5 LCLs carrying *FTSJ1* loss-of-function variants compared to the 4 LCLs from control individuals. The principal component analysis (PCA) from the different *FTSJ1* loss-of-function cell lines shows a high similarity and thus clusters on the PCA plot, while the wild type lines were more dispersed, possibly explained by their geographic origins (Figure S3A). The DESeq2 differential expression analysis showed statistically significant deregulation of 36 miRNAs when comparing *FTSJ1* mutants to control LCLs. 17 miRNA were up- and 19 down-regulated (Figures 3A, S3B and log2 FC and adjusted *p* values in Table S1). Importantly, as already reported in *Drosophila* (*Angelova and Dimitrova et al. 2020*), the global miRNA distribution was not drastically affected, thus ruling out general involvement of FTSJ1 in miRNA biogenesis.

**Figure 3.**
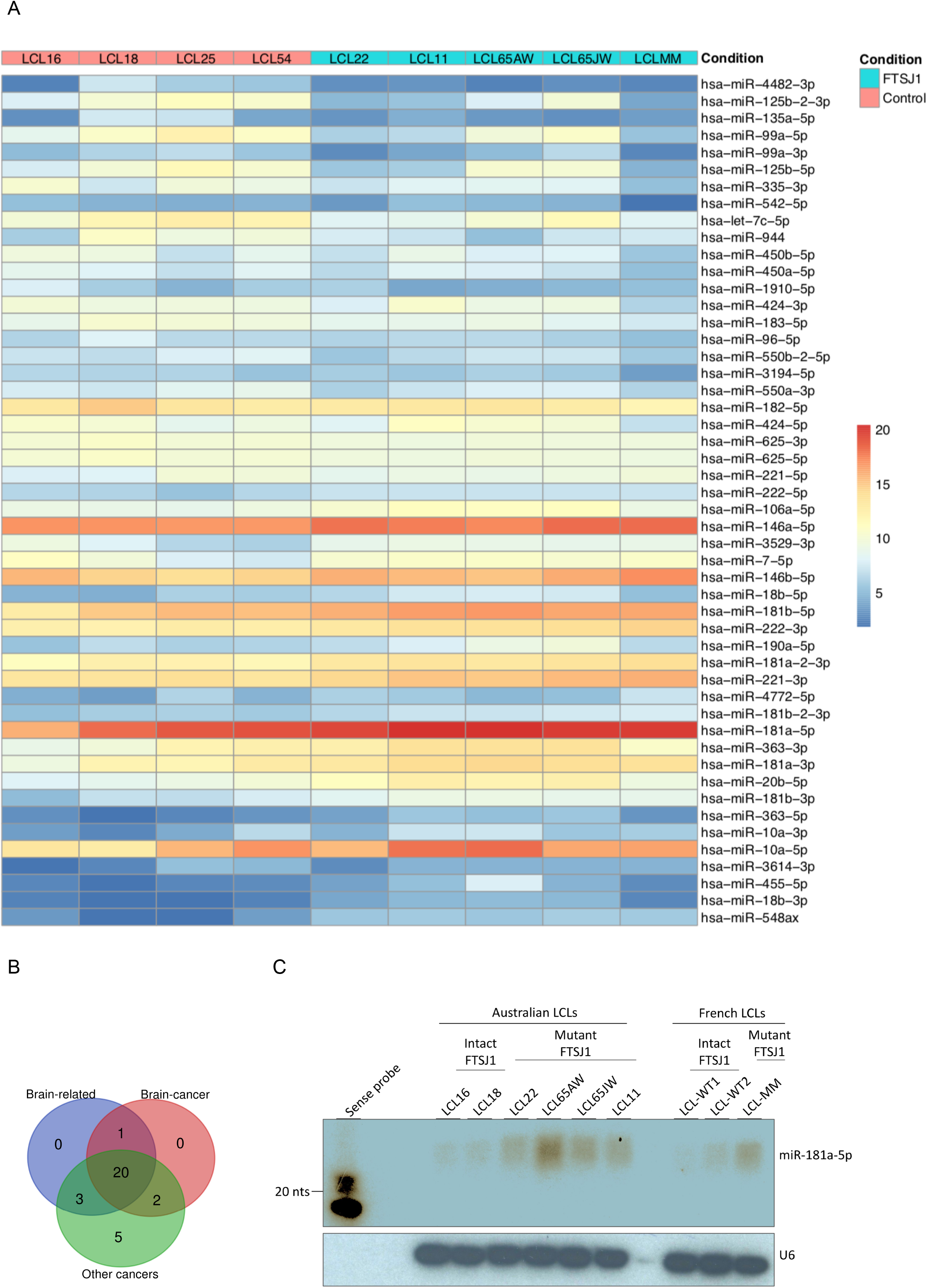
FTSJ1 loss of function leads to miRNAs deregulation in NSXLID affected individuals LCLs. **(A)** Heat map generated using the pheatmap package in R showing the 50 best deregulated miRNAs in p-values, and sorted fold change from most down-regulated (blue) to most up-regulated (red) are represented in two experimental conditions: FTSJ1 loss of function LCLs (blue turquoise) compared to controls LCLs (pink). Condition points to the FTSJ1 LCL status, WT (Control) or mutated for *FTSJ1* gene (FTSJ1). The data come from normalized and variance stabilizing transformed read counts using the DESeq2 package in R. **(B)** Bibliographic search (Table. 4) of the miRNAs deregulated in FTSJ1 loss of function LCLs reveals evidence for many of them as being implicated in cancers or brain development and brain diseases. The number of miRNAs related to brain, cancer and brain-cancer specifically are indicated respectively in the blue, green and red circle. The Venn diagram was generated by http://bioinformatics.psb.ugent.be/webtools/Venn/. **(C)** Northern blot analysis with ^32^P-labelled probe specific for hsa-miR-181a-5p confirms the upregulation of this miRNA in FTSJ1 loss of function condition already detected by small RNAseq analysis. A ^32^P-labelled probe specific for human U6 RNA was used to assess equal loading on the blot.

Next, we sought for possible links between the 36 significantly deregulated miRNAs in *FTSJ1* mutant cells and neuronal functions or neurodevelopmental disorders. Interestingly, 21 of these miRNAs were already identified in other small RNA-seq studies performed in the context of brain diseases such as epilepsy, Parkinson’s and Alzheimer’s diseases (Lau *et al*, 2013; Kretschmann *et al*, 2015; Ding *et al*, 2016; Roser *et al*, 2018). In addition, 29 of the deregulated miRNAs were linked to different types of cancers (Lund, 2010; Watahiki *et al*, 2011; Li *et al*, 2015; Khuu *et al*, 2016; Yang *et al*, 2017; Jiang *et al*, 2018), including 21 involved specifically in brain-related cancers, mostly in glioblastoma (Gillies & Lorimer, 2007; Shi *et al*, 2008; Lund, 2010; Conti *et al*, 2016) (Figure 3B and Table 4).

**Table 4.** Bibliographic search on miRNA deregulated in FTSJ1 loss-of-function LCL mutant cell line. The list shows for each miRNA if any link was found to brain development or brain-related diseases, also cancer and specifically to brain-cancers. The references are given for most of the miRNAs. The color code of the miRNA names indicates if they were found to be up- (red) or down-regulates (blue) in FTSJ1 mutant LCLs derived from NSXLID affected individuals compared to control LCLs derived from healthy individuals.

| miRNA | Brain related | Brain cancer related | Cancer related |
| --- | --- | --- | --- |
| hsa-miR-20b-5p | - | - | (Khuu *et al*, 2016) |
| hsa-miR-222-3p | (Lau *et al*, 2013)<br>(Kretschmann *et al*, 2015)<br>(Kan *et al*, 2012)<br>(Risbud & Porter, 2013) | (Gillies & Lorimer, 2007)<br>(Zhang *et al*, 2010) | - |
| hsa-miR-548ax | - | neuroblastoma for other<br>miR-548 family members | (Watahiki *et al*, 2011) (also<br>others cancers for other<br>miR-548 family members) |
| hsa-miR-125b-2-3p | yes | yes | yes |
| hsa-miR-221-3p | (Kretschmann *et al*, 2015)<br>(Kan *et al*, 2012)<br>(Risbud & Porter, 2013)<br>(Ding *et al*, 2016)<br>(Ma *et al*, 2016)<br>(Roser *et al*, 2018) | (see miR-222-3p) | (Fornari *et al*, 2008) |
| hsa-miR-335-3p | yes | yes | yes |
| hsa-miR-181b-2-3p | (see miR(181a-5p)) | (see miR(181a-5p)) | (see miR(181a-5p)) |
| hsa-miR-99a-5p | yes | yes | yes |
| hsa-miR-10a-5p | (Gui *et al*, 2015)<br>(Roser *et al*, 2018) | (Tehler *et al*, 2011)<br>(Lund, 2010) | (Tehler *et al*, 2011)<br>(Lund, 2010) |
| hsa-miR-181b-3p | (see miR(181a-5p)) | (see miR(181a-5p)) | (see miR(181a-5p)) |
| hsa-miR-106a-5p | yes | yes | yes |
| hsa-miR-181a-2-3p | (see miR(181a-5p)) | (see miR(181a-5p)) | (see miR(181a-5p)) |
| hsa-miR-146a-5p | yes | yes | yes |
| hsa-miR-4482-3p | - | - | - |
| hsa-miR-125b-5p | yes | yes | yes |
| hsa-miR-450b-5p | - | - | yes |
| hsa-miR-424-3p | yes | - | yes |
| hsa-miR-363-3p | (Lau *et al*, 2013)<br>(Kiyosawa *et al*, 2019) | (Conti *et al*, 2016)<br>(Qiao *et al*, 2013) | (Jiang *et al*, 2018)<br>(Ye *et al*, 2017)<br>(Hu *et al*, 2016)<br>(Wang *et al*, 2016)<br>(Karatas *et al*, 2016)<br>(Chapman *et al*, 2015)<br>(Zhang *et al*, 2016)<br>(Khuu *et al*, 2016) |
| hsa-let-7c-5p | - | - | - |
| hsa-miR-450a-5p | yes | - | yes |
| hsa-miR-18b-5p | - | - | - |
| hsa-miR-550a-3p | - | - | yes |
| hsa-miR-181a-5p | (Zhang *et al*, 2017)<br>(Ding *et al*, 2016)<br>(Roser *et al*, 2018) | (Shi *et al*, 2008) | (Yang *et al*, 2017)<br>(Li *et al*, 2015) |
| hsa-miR-550b-2-5p | - | - | yes |
| hsa-miR-181a-3p | (see miR(181a-5p)) | (see miR(181a-5p)) | (see miR(181a-5p)) |
| hsa-miR-181b-5p | (see miR(181a-5p)) | (see miR(181a-5p)) | (see miR(181a-5p)) |
| hsa-miR-183-5p | yes | yes | yes |
| hsa-miR-99a-3p | yes | yes | yes |
| hsa-miR-135a-5p | yes | yes | yes |
| hsa-miR-146b-5p | yes | yes | yes |
| hsa-miR-542-5p | - | yes | yes |
| hsa-miR-944 | yes | - | yes |
| hsa-miR-625-5p | - | - | - |
| hsa-miR-625-3p | - | - | - |
| hsa-miR-4772-5p | - | - | yes |
| hsa-miR-182-5p | yes | yes | yes |
| **Total #** | **24** | **23** | **30** |

To strengthen the small RNA-seq data, four hemizygous *FTSJ1* LCLs (control) and five LCLs mutants for *FTSJ1* were analysed by northern blotting with a specific probe complementary to *miRNA-181a-5p*. We selected this miRNA as it was highly upregulated in our small RNA-seq analysis and it was previously reported to be involved in vascular inflammation and atherosclerosis (Su *et al*, 2019), as well as expressed in neuronal cells in mammals (Dostie *et al*, 2003). One clear hybridization signal was observed in all *FTSJ1* mutant LCLs corresponding to mature *miRNA-181a-5p* (Figure 3C). In contrast, the 4 control LCLs show no or weak signal even after image over-exposure (Figures 3C). Together these results demonstrate that *FTSJ1* loss of function affects specifically the steady state levels of some miRNA and suggests that the deregulation of miRNA-mediated gene silencing observed in *FTSJ1* mutant LCLs was not caused by a global failure in miRNA biogenesis (Figures 3A, S3B and Table S1).

### FTSJ1 mutation perturbates the silencing activity of *miR-181a-5p* miRNA

As some of the FTSJ1 deregulated miRNAs and mRNAs were implicated in similar biological processes such as cancer and brain function, we wondered if there were some miRNA::mRNA pairs that could be involved in these commonly deregulated processes. Using miRNet 2.0 (Chang *et al*, 2020), we performed a bioinformatics cross-analysis of the small RNA-seq and mRNA-seq datasets. We found a subset of *FTSJ1*-deregulated miRNAs that were previously shown to modulate some of the FTSJ1 deregulated mRNAs. For instance, the *SPARC* mRNA is an experimentally confirmed target of *mir-10a-5p* (*Bryant et al, 2012; Wang et al, 2020*). This result thus suggests that *SPARC* mRNA downregulation observed in *FTSJ1* mutants may be due to its increased silencing by the upregulated *miR-10a-5p*. This cross-analysis also revealed that the *BTBD3* gene is potentially targeted by *miR-181a-5p (He et al, 2015)*, the two of which were upregulated in NSXLID affected individuals-derived LCLs (Figures 3A, 3C and Table 4), implicating a possible connection between them that differs from the canonical miRNA silencing pathway. LCL are known to be hardly transfectable (Nagayoshi *et al*, 2021), however *miR-181a-5p* and *BTBD3* are expressed similarly in HeLa cells (Figure S4A). Thus, by mimicking *miR-181a-5p* expression or repression, we show that *miR-181a-5p* silences *BTBD3* in HeLa cells (Figure S4B), strongly suggesting that *BTBD3* mRNA is a *bona fide* target of *miR-181a-5p*. Strikingly, in FTSJ1 mutant cells, the silencing activity of *miR-181a-5p* on *BTBD3* is compromised in both HeLa and LCL. Interestingly, despite the fact that 39 *ZNF* mRNAs were found potentially regulated by *miR-181a-5p* (Table 4 and (He *et al*, 2015)) and the over-representation of this miRNA in *FTSJ1* mutant (Figures 3A, 3C and Table S1), no evidence of miRNA regulation was yet found for *ZNF711*, a gene previously reported to be involved in the development of ID (van der Werf *et al*, 2017).

### FTSJ1 is involved in human neuronal morphology during development

The loss of *FTSJ1* in humans gives rise to ID, yet the underlying mechanism is still unclear. Both neuronal morphology (Chen *et al*, 2020) and behaviour (Jensen *et al*, 2019) have been reported in patients affected by a wide range of ID disorders, with a variety of genetic etiologies and their corresponding mouse models. To address whether loss of human *FTSJ1* also affects neuronal morphology, we altered FTSJ1 activity using 2,6-Diaminopurine (DAP) (Palma & Lejeune, 2021; Trzaska *et al*, 2020b) in human Neural Progenitor Cells (NPC). DAP is a recently discovered drug that binds to FTSJ1 and inhibits its methylase activity (Palma & Lejeune, 2021; Trzaska *et al*, 2020b). Immunostainings were performed for Sox2, a transcription factor expressed in NPCs, and Doublecortin (DCX), an associated microtubule protein expressed in differentiating NPCs or immature neurons, reflecting neurogenesis. Importantly, the DAP treatment did not significantly affect the differentiation of the NPCs (DCX-) to immature neurons (DCX+) (Figure 4A). This is in agreement with previous reports showing the absence of severe brain morphological defects in mice mutated for FTSJ1 (Jensen *et al*, 2019; Nagayoshi *et al*, 2021). However DCX positive cells treated with 100μM DAP showed a 25% increase in the number of interstitial protrusions, likely filopodia, on their neurites compared to the smoother appearance of the neurites of untreated control cells (Figures 4B and 4C). These spines’ morphological defects on DAP treated DCX+ cells are reminiscent of those observed on mature neurons from mutant mice of the Fragile X mental retardation protein (FMRP) (Braun & Segal, 2000), as well as from human patients’ brains that suffer from the fragile X syndrome. Furthermore similar findings were recently reported in mice brains mutated for *FTSJ1* (Nagayoshi *et al*, 2021), suggesting that this is a conserved phenotypic consequence of the loss of FTSJ1.

**Figure 4.**
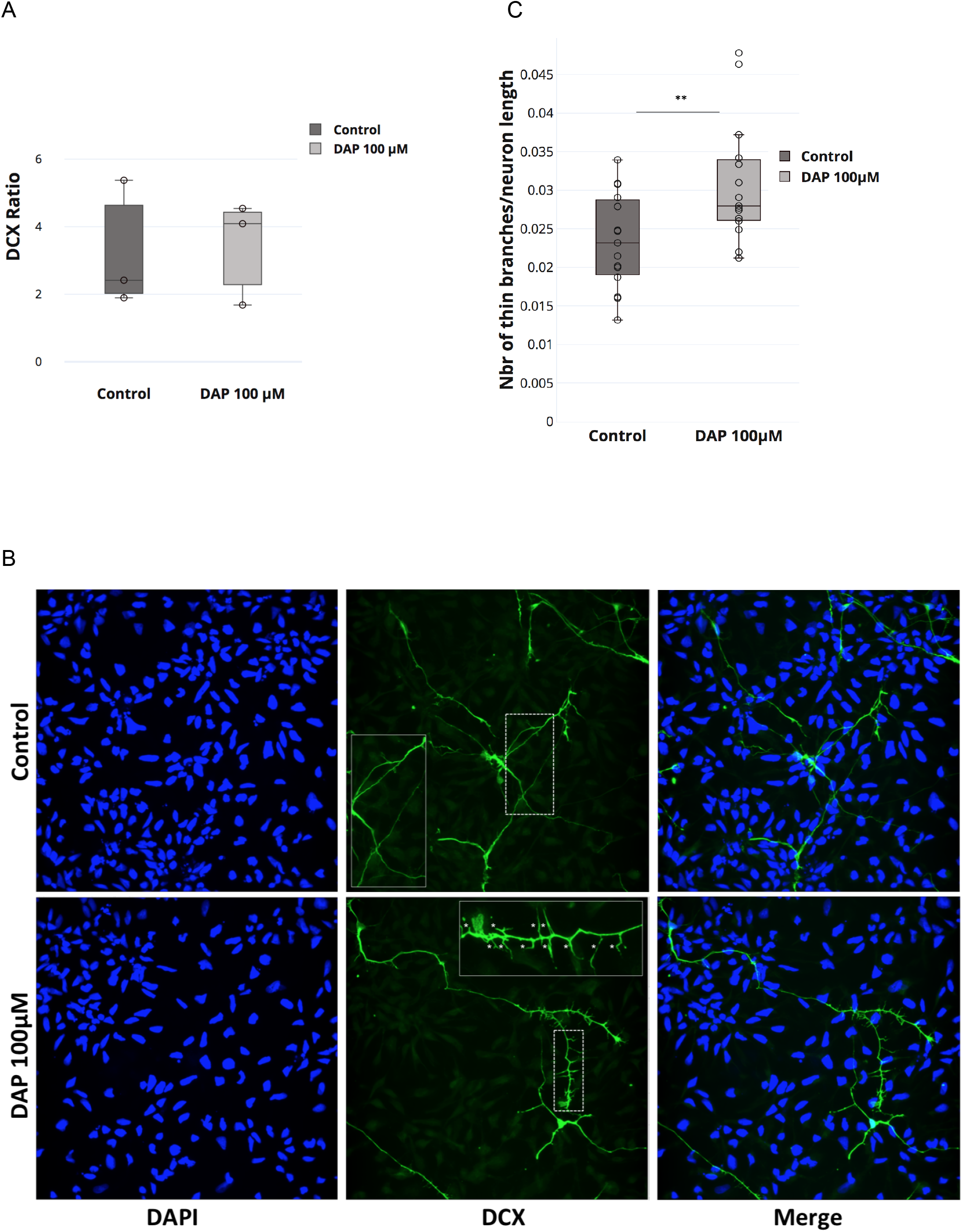
FTSJ1 depletion affects human neurons’ neurite spines morphology. **(A)** DAP induced FTSJ1 inhibition does not affect human NPC to immature neuron differentiation. Immunostainings for DCX and SOX2 were performed on human iPSCs derived NPCs either treated with 100μM DAP or equal volume of H_2_O for 24h. Cells were numbered on microscopy acquisitions, and the ratio of DCX expressing cells over total cell number was calculated and expressed in fold change. Error bars represent standard deviation of three independent experiments; n.s: not significant (over 1400 cells numbered for a single experiment). (**B)**. **Lower panel**: Human NPCs inhibited for FTSJ1 with 100 μM DAP for 24h (DAP 100μM) present an increased number of neurite spines during NPC to immature neuron differentiation. DCX protein expressed in immature neurons is marked in green (DCX). Dashed white line represents the *zoom-in* zone depicted in the top right corner with a continuous white line. White stars (*) in the magnified inset point to the fine spine neurites. **Upper panel**: Untreated NPCs (Control). Nuclear staining was performed using DAPI depicted in blue (DAPI). **(C)** Quantification of thin spines of DCX positive cells (Figure 4B above). Thin projections were numbered and normalized over the total length of the immature neurons as traced and measured by SNT (Fiji plugin). Quantifications were carried out on 5 acquisitions for each experiment (Control and DAP 100 μM) (>40 branches/acquisition on average). Aggregate of 3 independent experiments. Wilcoxon Mann-Whitney’s test **P= 0,0098.

### *Drosophila* FTSJ1 ortholog is involved in neuronal morphology during development

To further address whether the control of neuron morphology by FTSJ1 is a conserved feature across evolution we dissected the neuromuscular junctions (NMJs) of *Drosophila* larvae carrying mutations in the orthologs of the *FTSJ1* gene as well as larvae fed with DAP (Palma & Lejeune, 2021; Trzaska *et al*, 2020b). Examination of the NMJs in Trm7_32 and Trm7_34 double homozygous mutant larvae or larvae fed with DAP revealed a significant synaptic overgrowth when compared to control larvae (Figure 5). Furthermore, as observed for the human NPC treated with DAP (Figures 4B and 4C), the neurite branching was strongly increased in both, double mutant and fed treated larvae (Figure 5). However the overall length of the axons was not significantly altered. These results indicate that *Drosophila* FTSJ1s, alike human FTSJ1, control neuronal morphology.

**Figure 5.**
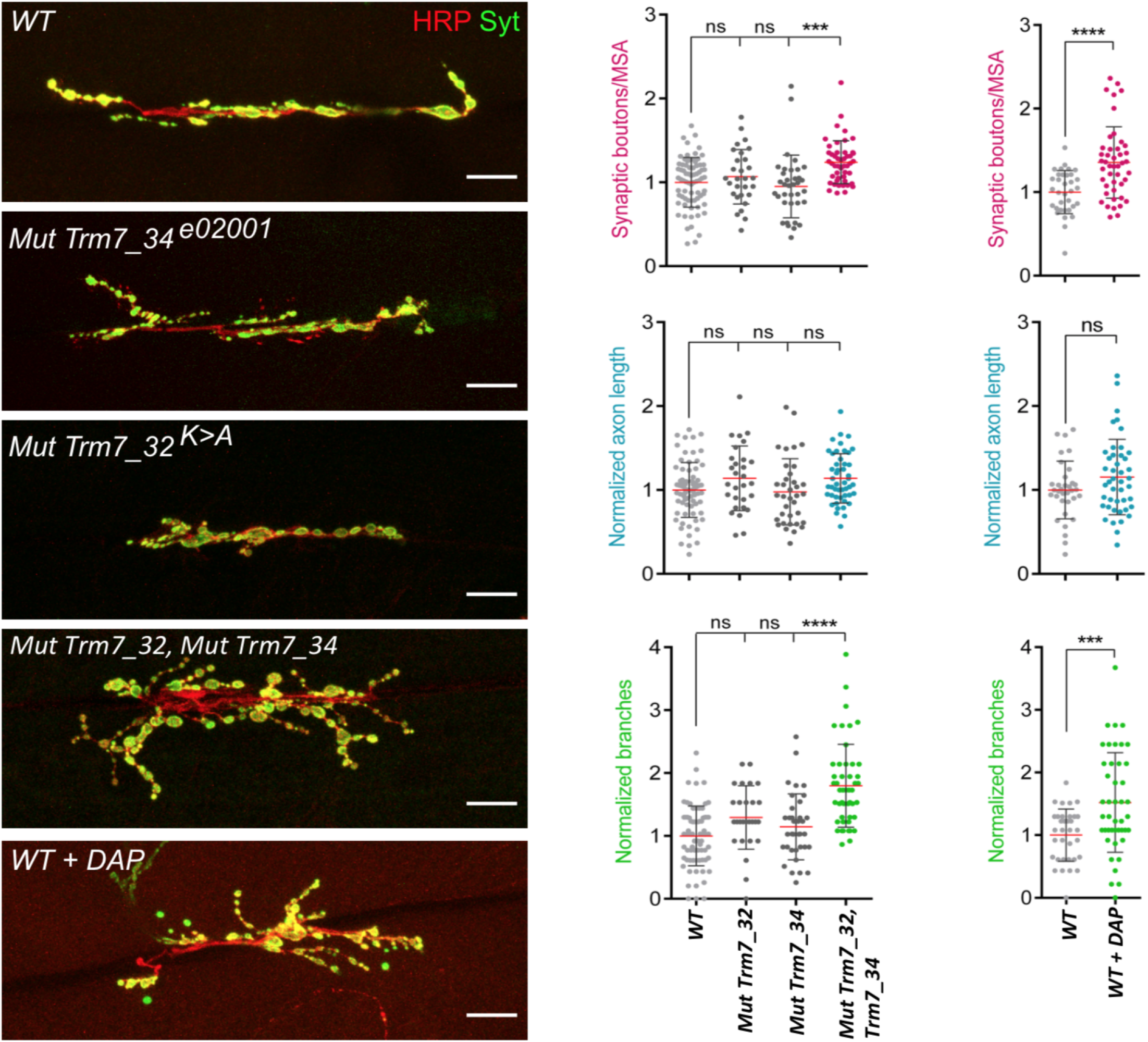
FTSJ1 dependent Nm regulates axonal morphlogy in the *Drosophila* nervous system. **Left panel:** Representative confocal images of muscle-6/7 NMJ synapses of larval abdominal hemisegments A2–A3 for the indicated genotypes labelled with anti-synaptotagmin (green) and HRP (red) to reveal the synaptic vesicles and the neuronal membrane. Scale bar: 20 μm. **Right panel:** Quantification of normalized bouton number (total number of boutons/muscle surface area (MSA) (μm^2^ × 1,000)) (**top**), normalized axon length (**middle**) and normalized branching (**bottom**) of NMJ 6/7 in A2–A3 of the indicated genotypes. Bars show mean ± s.e.m. Multiple comparisons were performed using one-way ANOVA with a post hoc Sidak–Bonferroni correction. (ns. = not significant; *P < 0.05; ***P < 0.001; ****P < 0.0001). Numbers of replicated neurons (n) are: 74 for WT; 36 for Trm7_32; 29 for Trm7_34; 48 for Trm7_32, Trm7_34 and 34 for WT untreated and 45 for WT treated with DAP. *Canton-S* larvae were used as wild-type control.

### Reward learning requires FTSJ1 activity in *Drosophila*

*FTSJ1* loss of function affected individuals suffer from significant limitations both in intellectual functioning and in adaptive behaviour. Similar phenotypes including impaired learning and memory capacity were recently observed in *FTSJ1* KO mice that also present a reduced body weight and bone mass, as well as altered energy metabolism (Jensen *et al*, 2019; Nagayoshi *et al*, 2021). In flies, we recently showed that the loss of FTSJ1 orthologs causes reduced lifespan and body weight, as well as locomotion defects (Angelova and Dimitrova et al. 2020).

To address whether fly memory was also altered in these mutants we applied the appetitive conditioning assay. We found that short-term memory (STM) of single *Trm7_34* or *Trm7_32* and double *Trm7_34;Trm7_32* heterozygous mutant flies was indistinguishable from that of wild-type controls (Figure 6A). However, long-term memory (LTM) was significantly impaired in all of these three mutant combinations (Figure 6B). Importantly, naive heterozygous mutants flies detected sugar properly and behave normally when exposed to repellent odors used in the olfactory memory assay (Figures 6C and 6D), suggesting that the LTM defect was not due to a confounding alteration of sensory abilities. Thus, these results indicate that the *Drosophila* FTSJ1 ortholog Trm7_34 and Trm7_32 has a specific function in LTM, and importantly demonstrate clearly that both tRNA Nm32 and Nm34 modifications have function in long term memory.

**Figure 6.**
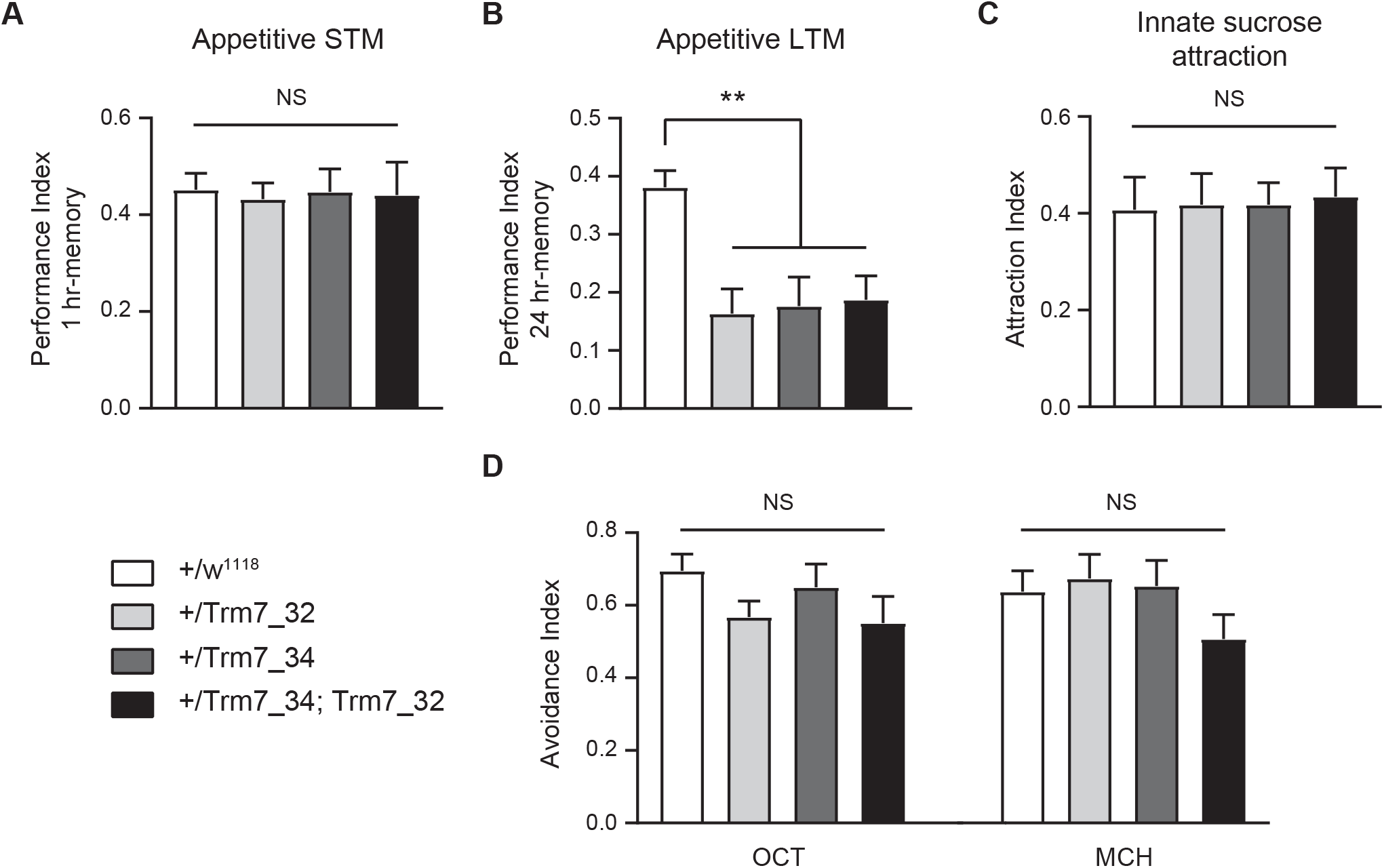
FTSJ1 *Drosophila* FTSJ1 ortolog *Trm7_34* mutants are defective for appetitive long-term memory. Behavioral performances are reported as mean +/−SEM. Statistical significance was tested with a one-way ANOVA followed by Tukey posthoc pairwise comparisons. Asterisks on the barplots indicate the level of significance of the pairwise comparison with control. The *p*-value indicated in the legend corresponds to the output of the ANOVA. **(A)** Flies were starved on mineral water for 21 hrs and then trained with an appetitive associative olfactory learning protocol (odor paired with sucrose ingestion). Short-term memory (STM) performance was measured 1 hr after learning. The STM score of flies heterozygous mutants for Trm7_32 (+/Trm7_32), Trm7_34 (+/Trm7_34), and double heterozygous Trm7_32; Trm7_34 (Trm7_32;Trm7_34) were not different from their genotypic controls (+/w^1118^) (*n* = 12 per condition; *p* = 0.99). **(B)** Flies were starved on mineral water for 21 hrs and then trained with an appetitive associative olfactory learning protocol (odor paired with sucrose ingestion). Long-Term Memory (LTM) performance was measured 24 hrs after learning. The LTM score of flies heterozygous mutants for Trm7_32 (+/Trm7_32), Trm7_34 (+/Trm7_34), and double heterozygous Trm7_32; Trm7_34 (Trm7_32;Trm7_34) were severely impaired as compared to their genotypic controls (+/w^1118^) (*n* = 16–19 per condition; *p* = 0.0007). **(C)** Flies were starved on mineral water for 21 hrs, and their attraction to sucrose was then measured. The innate sucrose preference of flies heterozygous mutants for Trm7_32 (+/Trm7_32), Trm7_34 (+/Trm7_34), and double heterozygous Trm7_32; Trm7_34 (Trm7_32;Trm7_34) were not different from their genotypic controls (+/w^1118^) (*n* =14 per condition; *p* = 0.99). **(D)** Flies were starved on mineral water for 21 hrs, and their avoidance to the odorants used in the olfactory memory assays, 3-octanol (OCT) and 4-methylcyclohexanol (MCH) was then measured. The innate odor avoidance of flies heterozygous mutants for Trm7_32 (+/Trm7_32), Trm7_34 (+/Trm7_34), and double heterozygous Trm7_32; Trm7_34 (Trm7_32;Trm7_34) were not different from their genotypic controls (+/w^1118^) (*n* = 10 per condition; OCT: *p* = 0.26; MCH: *p* = 0.28).

## DISCUSSION

In this study, we characterised at the molecular and cellular levels the effect of *FTSJ1* loss of function in human cells. We used the innovative RiboMethSeq method to analyse the Nm status from five patients carrying distinct loss of *FTSJ1* functions, which led us to the identification of new human FTSJ1 tRNA targets. Furthermore we identify specific transcripts and miRNA that are misregulated in the absence of FTSJ1, that may contribute to the FTSJ1 pathologies, and suggest potential cross-regulation among them. Lastly we show for the first time that the lack of FTSJ1 alters the morphology of human neurons, a phenotype that is conserved in *Drosophila* and is associated with long term memory deficit.

The power of the RiboMethSeq approach is that it allows to analyse the Nm status of the totality of transcribed tRNA species and not only selected tRNAs based on the prior but incomplete knowledge of FTSJ1 targets. Furthermore, this approach covers the whole tRNA-ome and thus can identify variations in Nm at the single nucleotide resolution, which is very useful to distinguish tRNA isoacceptors for instance that differ by only few nucleotides. Our results from the RiboMethSeq performed on patient and control LCLs confirmed the already known human tRNA targets of FTSJ1. For instance, Cm_32_ and Cm_34_ of tRNA^Trp(CCA)^ as well as position 34 in tRNA^Phe(GAA)^ and tRNA^Leu(CAG)^ were validated by our approach. Only Cm_32_ of tRNA^Phe(GAA)^, which is a well-known target of FTSJ1, could not be validated at the first glance. The analysis of this position is challenging due to low read numbers necessary for its quantification. This is the result of two confounding factors. On one hand the calculation of MethScores (Figure 1A) is based on the two neighbouring nucleotides (Marchand *et al*, 2016) Since FTSJ1 deposits Nm at both 32 and 34 positions in tRNA^Phe^, the calculated MethScore at position 32 is affected when position 34 of the same tRNA is also Nm modified. Second, we previously reported that tRNA^Phe(GAA)^ ACL positions are challenging to detect due to the specific hyper-modification on position 37 of tRNA^Phe^ (Angelova and Dimitrova et al. 2020). Indeed, o2yW_37_/m^1^G_37_ impairs reverse transcription thereby reducing the number of cDNAs spanning the ACL. Nevertheless, deeper visual inspection of the raw reads profile shows that Nm at position 32 was indeed lost in *FTSJ1* mutated cells when compared to control LCL (Figure S1D), confirming the previous reports.

Importantly, we confirmed recent (tRNA^Arg(UCG)^ and tRNA^Gln(CUG)^) and identified novel (tRNA^Gly(CCC)^, tRNA^Leu(UAA)^, tRNA^Pro^, and tRNA^Cys(GCA)^, Table 2) tRNA targets for human FTSJ1. In the case of tRNA^Arg(UCG)^, we confirmed not only a new target for FTSJ1, but also a modification which was not previously reported in modomics but only recently in HEK293 FTSJ1 CRISPR mutant (Li *et al*, 2020a). Indeed C_32_ is known to be m^3^C and not Nm modified for the two other isoacceptors (tRNA^Arg(CCU)^ and tRNA^Arg(UCU)^) (Boccaletto *et al*, 2018). Similarly, there was no evidence for a human Cm_32_ tRNA^Gln(CUG)^ and only the other isoacceptor tRNA^Gln(UUG)^ was reported in Modomics as 2’-O-methylated at C_32_. Still, Cm_32_ on tRNA^Gln(CUG)^ was recently discovered as a target of *Drosophila* Trm7_32 (Angelova and Dimitrova et al. 2020). Among the newly uncovered FTSJ1 targets in this study, Um_32_ tRNA^Gly(CCC)^ was the only one that has been reported in Modomics, however the enzyme responsible for this modification was yet unknown. Our results demonstrate that FTSJ1 is the dedicated human Nm-Mtase that installs Um_32_/Cm_32_ and Cm_34_/Um_34_/Gm_34_ residues on human tRNAs.

Our transcriptomic analysis also highlighted novel transcripts and miRNA targets that may play important roles in the development of the diseases. For instance we found 36 differentially expressed miRNAs, most of which were already associated with brain diseases and functioning and/or cancer development. Strikingly, the most prevalent associated cancer types were the ones related to the brain tissues. Consistently with the post-transcription regulation role of miRNA, we also found through mRNA-seq an enrichment of brain morphogenesis-related mRNAs differentially expressed in *FTSJ1* loss of function when compared to control LCLs. Interestingly, a cross-analysis of these two RNA sequencing experiments revealed potential miRNA::target mRNA couples among the deregulated RNA populations. This is indicative of possible miRNA silencing changes in the absence of FTSJ1, similarly to what we report earlier in *Drosophila FTSJ1* mutant orthologs. The predicted miRNA::mRNA couples need to be further validated individually in neuronal tissues, although their report from *miRnet* database (Chang *et al*, 2020) already includes experimental evidence on the miRNA::mRNA regulation, particularly for *BTBD3* and *SPARC* mRNAs (Bryant *et al*, 2012; Wang *et al*, 2020; He *et al*, 2015). In addition to the reported prediction (He *et al*, 2015), we show that *BTBD3* is a *bona fide miR-181a-5p* target. Surprisingly, both *BTBD3* and *miR-181a-5p* were up-regulated in FTSJ1 depleted patient cells. (Angelova and Dimitrova et al. 2020)suggest that Nm-MTases genes could act upstream of small RNA biogenesis and function through transcriptional downregulation of Argonaute mRNA in *Drosophila* FTSJ1 mutants (Angelova and Dimitrova et al. 2020) and in human cells (not shown). On the other hand, tRNA fragments (tRF) abundance seen in FTSJ1 mutant fly (Angelova and Dimitrova et al. 2020) and mice (Nagayoshi *et al*, 2021) can associate with Dicer, Argonaute and Piwi proteins, thus affecting their silencing function. Such tRF-mediated titration of proteins away from canonical substrates has been previously reported in *Drosophila* and human cell lines (Durdevic *et al*, 2013; Goodarzi *et al*, 2015).

Affected individuals carrying mutations in *FTSJ1* suffer from ID (Guy *et al*, 2015; Freude *et al*, 2004; Ramser *et al*, 2004) but the mechanism underlying this pathology has remained elusive. A recent report from Nagayoshi et al. added some insight by showing that *Ftsj1* loss of function in mice provoke dendritic spine overgrowth at hippocampus and cortex neurons (Nagayoshi *et al*, 2021), suggesting that a similar alteration of neuron morphology may exist in human patients, which might impair their functioning. Indeed we observed long, thin protrusion in human neurons affected for FTSJ1 activity. These protrusions are very similar in size and shape to the dendritic spines observed in hippocampus and cortex neurons of *Ftsj1* loss of function mice (Nagayoshi *et al*, 2021). A similar observation was also described earlier for *FMRP* mutant mice (Braun & Segal, 2000) and *FMRP* human affected individuals’ brains suffering from ID (Irwin *et al*, 2000). More examples of improper neuron morphology and in particular spine immaturity were found in additional gene loss of functions causative of ID (Levenga & Willemsen, 2012). This suggests that the lack of proper neuronal morphology may be a common feature of ID. More work will be required to address how these changes in spine arborization occur in the absence of FTSJ1 and how this translate into the disease. Interestingly in this study we found that *BTBD3* mRNA is significantly upregulated in FTSJ1 mutated LCLs. Since BTBD3 controls dendrite orientation in mammalian cortical neurons (Matsui *et al*, 2013) it will be an interesting target to further characterize in the context of FTSJ1 ID pathology.

A synaptic overgrowth was also observed in *Drosophila*, indicating that this function of FTSJ1 is conserved across evolution. In addition we found that the long term memory but not the short term was significantly altered in the absence of FTSJ1 in flies. This is consistent with the learning deficits observed in mice and humans. In contrast to Human FTSJ1 and the yeast ortholog TRM7, *Drosophila* uses two distinct paralogs to methylate positions 32 and 34, respectively, on tRNAs ACL. Interestingly, we found that the lack of both, Trm7_34 and Trm7_32 had an effect on long term memory, suggesting that the methylation at wobble position 34 and 32 are critical for this function. However, the lack of both modifications (as in mammals *Ftsj1* mutant) is not cumulative regarding the memory deficit (Figure. 6). This last observation is strongly supported by the affected human individual that harbours a missense variant (p.Ala26Pro, LCL22 in this study), resulting in loss of Gm_34_, but not of Cm_32_ in human tRNA^Phe^ (Guy *et al*, 2015). Further studies should aim to understand how the loss of methylation at these ACL positions affects the learning and memory functions.

The heterogeneity of ID makes it extremely challenging for genetic and clinical diagnosis (Ilyas *et al*, 2020). Our RiboMethSeq and transcriptomics approaches performed on NSXLID affected individuals have with high confidence extended the panel of FTSJ1’s targets. Since our investigation was carried out on LCLs derived from the blood of affected individuals, our resource provides potential new biomarkers for diagnosis of FTSJ1-related ID in the future. For instance, *miR-181a-5p*, which is detected only in patient derived blood cells, constitutes already a good candidate for such purpose. Therefore our study highlights the usefulness of companion diagnostics in clinical settings, in addition to exome sequencing, for potential discovery of prognostic markers of complex diseases.

## DESCRIPTION OF SUPPLEMENTAL DATA

Supplemental Data include 9 figures and 1 table.

## ACKNOWLEDGMENTS

We thank the patients and their families for their participation in the study. We also thank Christelle Thibaut-Charpentier from the GenomEast sequencing platform in Strasbourg, a member of the ‘France Génomique’ consortium (ANR-10-INBS-0009), the Institut de Génétique Médicale d’Alsace for their technical support and Myriam Bronner from Nancy University Hospital for the establishment of the LCL-MM line. We thank Johann Schor and Laura Guédon for their help with behavioral experiments. Human NPC work was carried out at ICM’s CELIS core facility and NPC imaging at the ICM Quant facility. We thank Dr. Bernard Moss (NIAID/NIH) for providing FTSJ1 KO HeLa cells. We thank the members of the TErBio laboratory for helpful discussions and reading of the manuscript. C.C. received financial support from the CNRS, Sorbonne Université (Emergence 2021_RNA-Mod-Diag), the Fondation Maladies Rares (Genomics-2018 #11809 & Genomics-2020 #12824), the IBPS-2020 Action Incitative, the Ligue National contre le cancer Île de France (RS21/76-29) and ANR (ANR-21-CE12-0022-01 #BiopiC). Work in the B.A.H. lab was supported by the Investissements d’Avenir program (ANR-10-IAIHU-06), Paris Brain Institute-ICM core funding, the Roger De Spoelberch Foundation Prize and a grant from the Neuro-Glia Foundation. Research in the laboratory of J.-Y.R. is supported by University of Lausanne and the Deutsche Forschungsgemeinschaft RO 4681/9-1, RO4681/12-1 and RO4681/13-1. D.G.D. and M.B. have PhD fellowships from the Ministère de la Recherche et de l’Enseignement Supérieur at the doctoral school Complexité du Vivant (ED515). We also thank the Fondation ARC pour la Recherche sur le Cancer and the FRM (Fondation pour la Recherche Médicale) for funding support to D.G.D., M.T.A and M.B. 4th years PhD; ‘Réseau André Picard’, the ‘Société Française de Génétique’ (to M.B., M.T.A and D.G.D.) and COST action ‘EPITRAN’ CA16120 (to Y.M., J.Y.R., V.M., D.G.D., M.B., M.T.A and C.C.) for travelling and training fellowships.

## DATA AND CODE AVAILABILITY

The RNA sequencing and small RNA sequencing data discussed in this publication are deposited and fully accessible upon request during the reviewing process, either in NCBI’s Gene Expression Omnibus accessible through GEO Series accession number GSE179384 for small RNAseq or at the European Nucleotide Archive (ENA) at EMBL-EBI under accession number PRJEB46400 for the RiboMethSeq and PRJEB46399 for RNA seq.

