## Supplementary Figures and TableS1 for "The ribose methylation enzyme FTSJ1 has a conserved role in neuron morphology and learning performance"

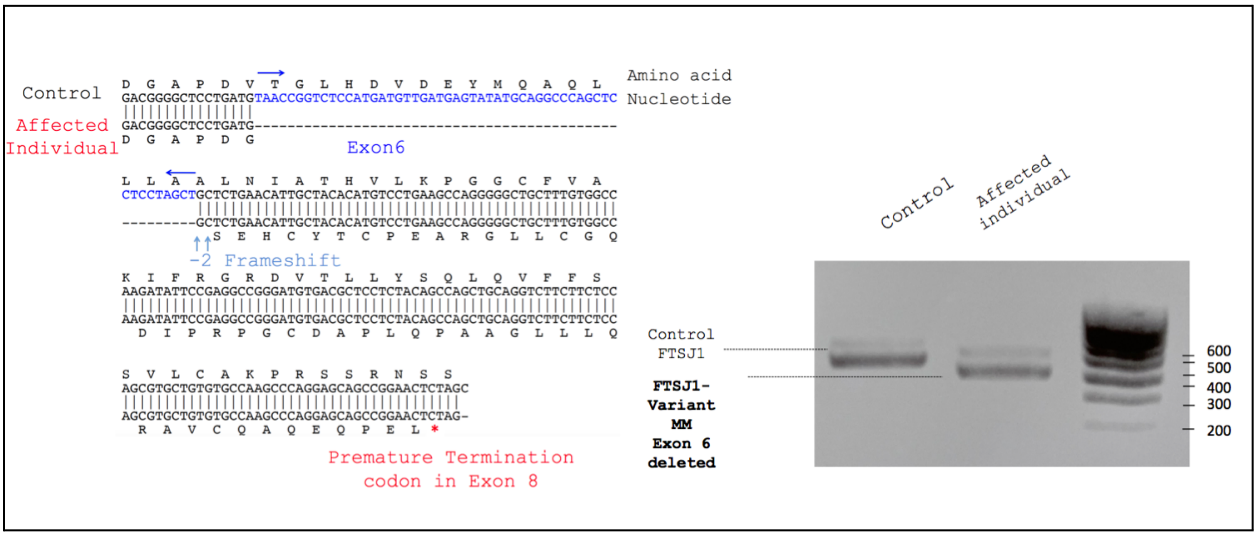


**Figure S1A: Characterization of the FTSJ1 transcript variant MM.** FTSJ1 transcripts were amplified from total RNA by RT-PCR of a control (LCL25) and the affected individual LCL MM harbouring a splice site mutation predicted to cause the skipping of exon 6 (see Mat & Met section). PCR products were run in 1% Agarose Gel + 0,5X TAE buffer. Predictably, a size shift is observed between the amplified products in the control as compared to the MM variant (Right Panel). Sanger sequencing confirmed skipping of the entire exon 6, thus disrupting the reading frame and appearance of a premature termination codon in Exon 8 (Left Panel, see detail in the Mat & Met section).

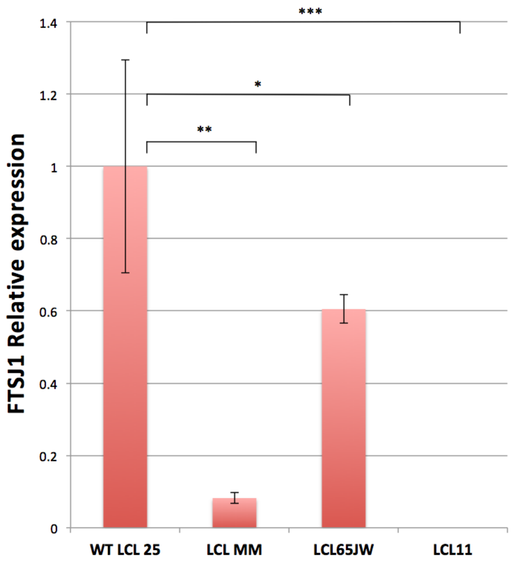


**Figure S1B. FTSJ1 mRNA relative abundance in patient LCLs.** FTSJ1 mRNA is significantly reduced in LCL MM compared to a control LCL. Relative abundance of mRNA is quantified by RT-qPCR on total RNAs from each indicated cell line. Ratios are expressed in fold change of starting quantities of FTSJ1/GAPDH. Error bars represent the standard deviation between four independent biological samples. P values are indicated with stars *p=0,04, **p=0,006, ***p=0,0008 (Paired Student's *t* test).

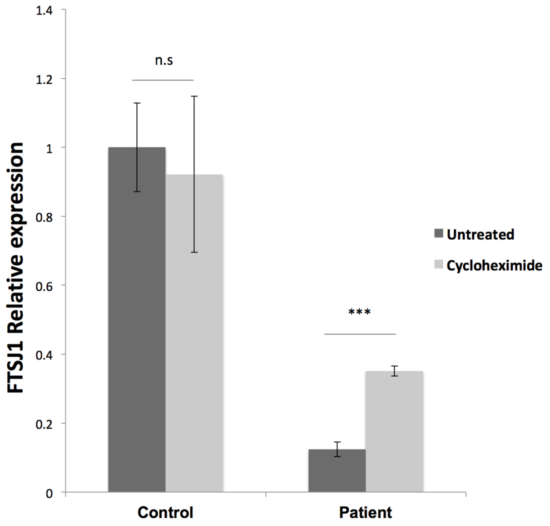


**Figure S1C. FTSJ1 mRNA targeted for degradation by Nonsense-mediated mRNA Decay (NMD) in LCL-MM.** FTSJ1 mRNA is significantly downregulated in affected patient cells (LCL-MM) when compared to control cells (LCL-25) (Figure S1B above). Inhibition of translation by treatment of LCLs with 100 µg ml–1 cycloheximide rescued FTSJ1 mRNA from NMD, resulting in a 3 fold increase as analysed by RT-qPCR. Starting quantity values were normalized against GAPDH and expressed in fold change. ***P=0,0008 (Mann Whitney test). Error bars indicate s.d.N=6.

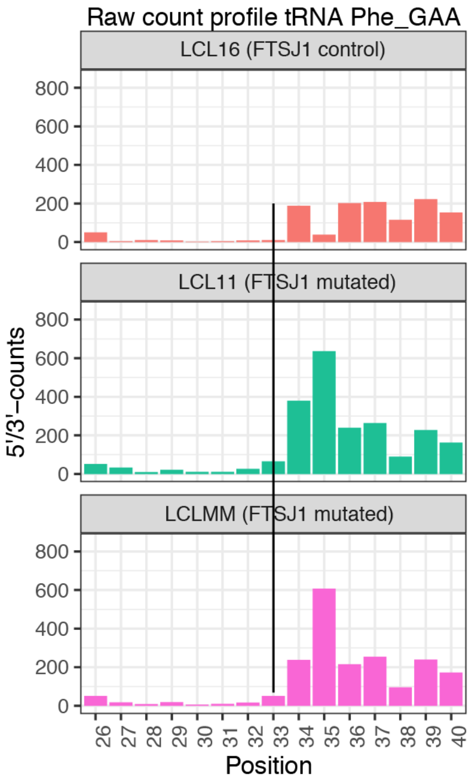


**Figure S1D. FTSJ1 targets human tRNA$^{Phe(GAA)}$ at position 32**. Related to Figure 1B. RiboMethSeq analysis of tRNA$^{Phe(GAA)}$ modification at positions $Cm_{32}$ and $Gm_{34}$ Alkaline fragmentation-based RiboMethSeq was performed on total RNAs extracted from indicated LCLs hemizygous mutants for *FTSJ1* (LCL11 and LCLMM) and control *FTSJ1* (non-mutated: LCL16*)* as indicated. For better visualization, raw read counts are presented in a non-normalized fashion (5'/3'-counts, raw count profil). The positions of interest ($Cm_{32}$) in tRNA$^{Phe(GAA)}$ is indicated by a black line crossing the 3 graphs.

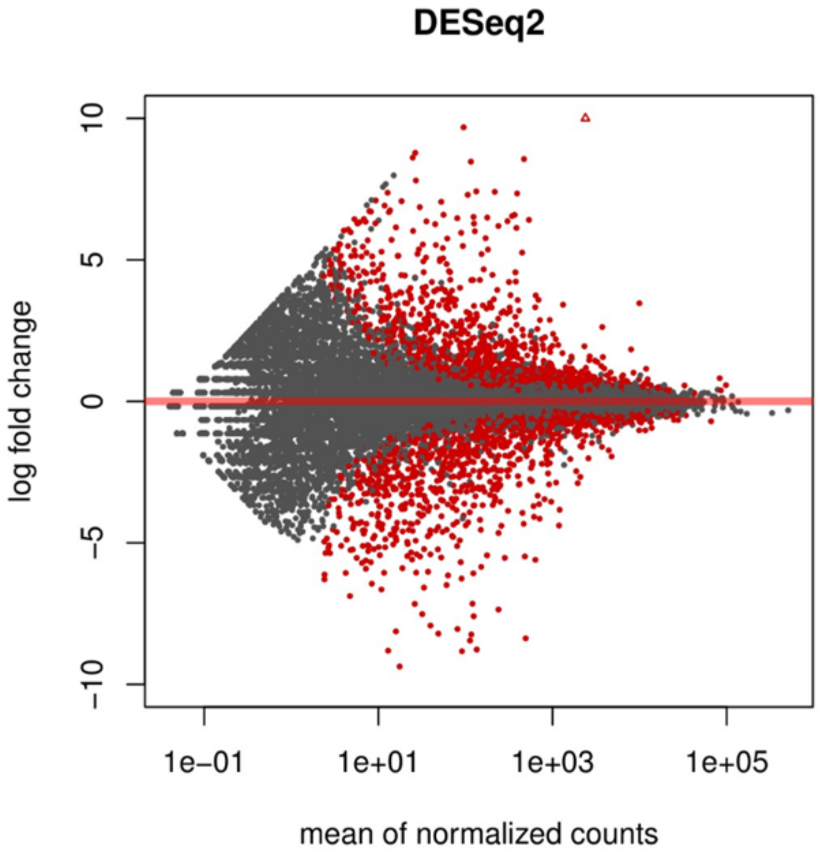


**Figure S2A. FTSJ1 loss of function leads to mRNAs deregulation in NSXLID affected individuals LCLs.** MAplot on data from sequencing of mRNAs showing multiple deregulated mRNAs.

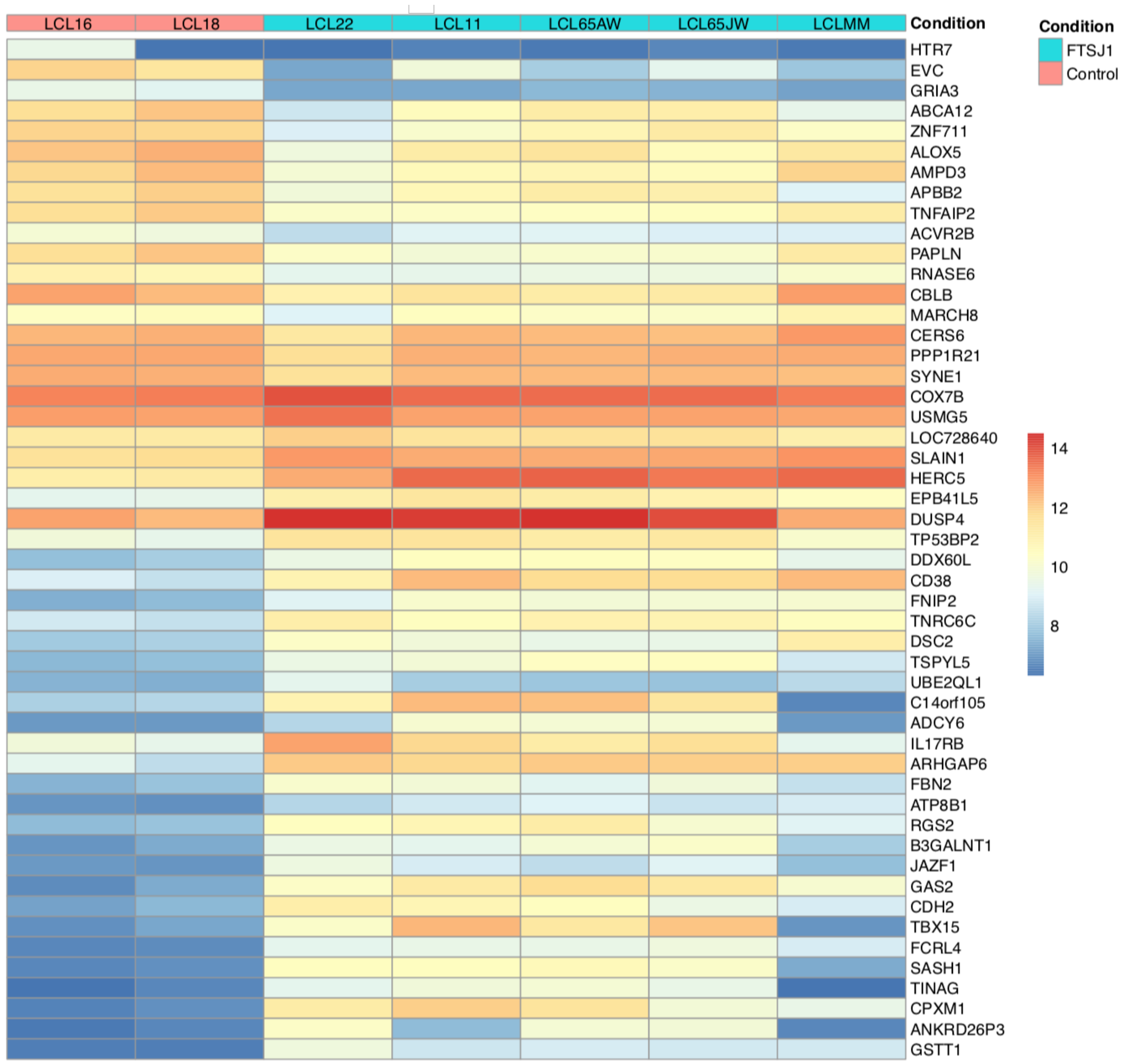


**Figure S2B. FTSJ1 loss of function leads to mRNAs deregulation in NSXLID affected individuals LCLs.** Heat map showing the top 50 deregulated mRNAs in p-values, and sorted fold change from most down-regulated to most up-regulated are represented in FTSJ1 loss of function LCLs compared to controls LCLs. Condition points to the FTSJ1 LCL status, WT (Control) or mutated for *FTSJ1* gene (FTSJ1). The data come from normalized and variance stabilizing transformed read counts using the DESeq2 package in R. Names of the LCLs are indicated in Condition lane.

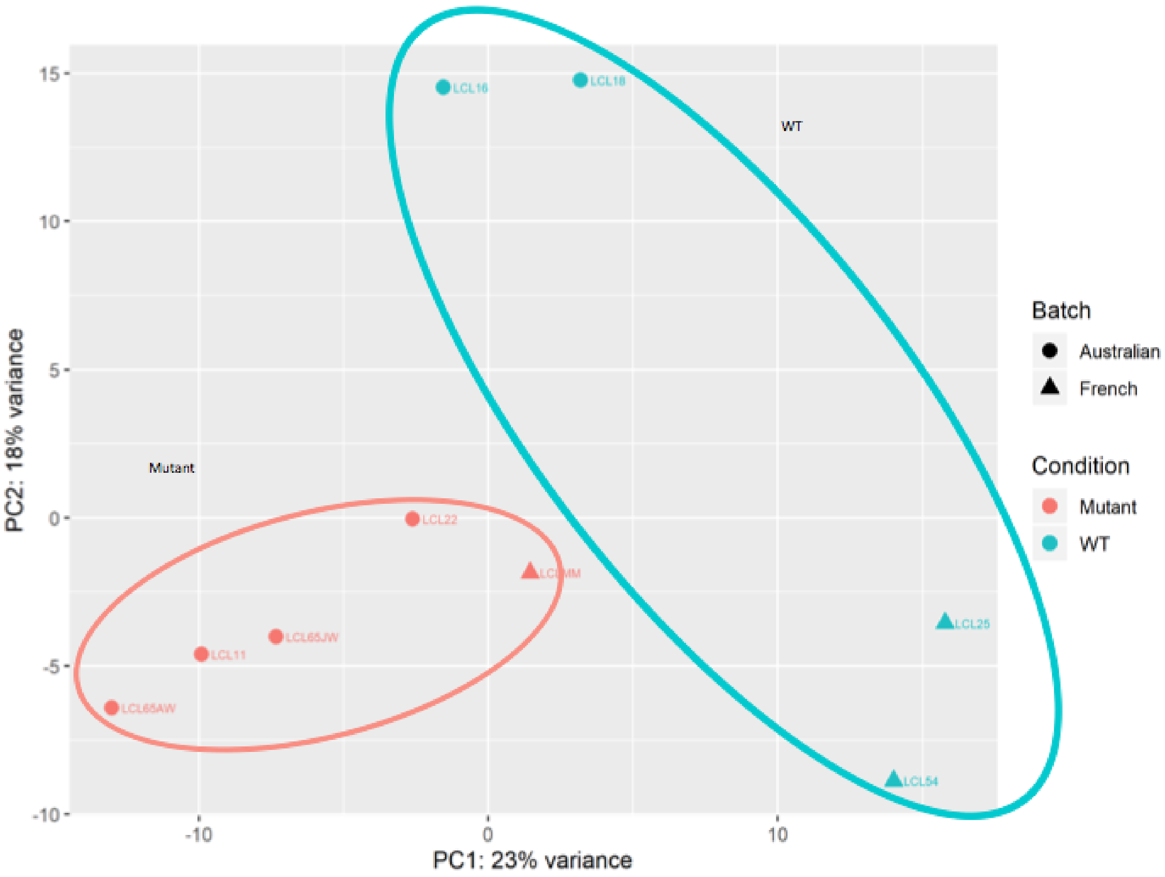


**Figure S3A. FTSJ1 loss of function leads to miRNAs deregulation in NSXLID affected individuals cells.** The principal component analysis (PCA) plot shows a well-defined cluster of all LCL lines lacking FTSJ1 function that is separated from the more dispersed cluster of control lines (LCL WT for FTSJ1: WT; LCL Mutant for FTSJ1: Mutant).

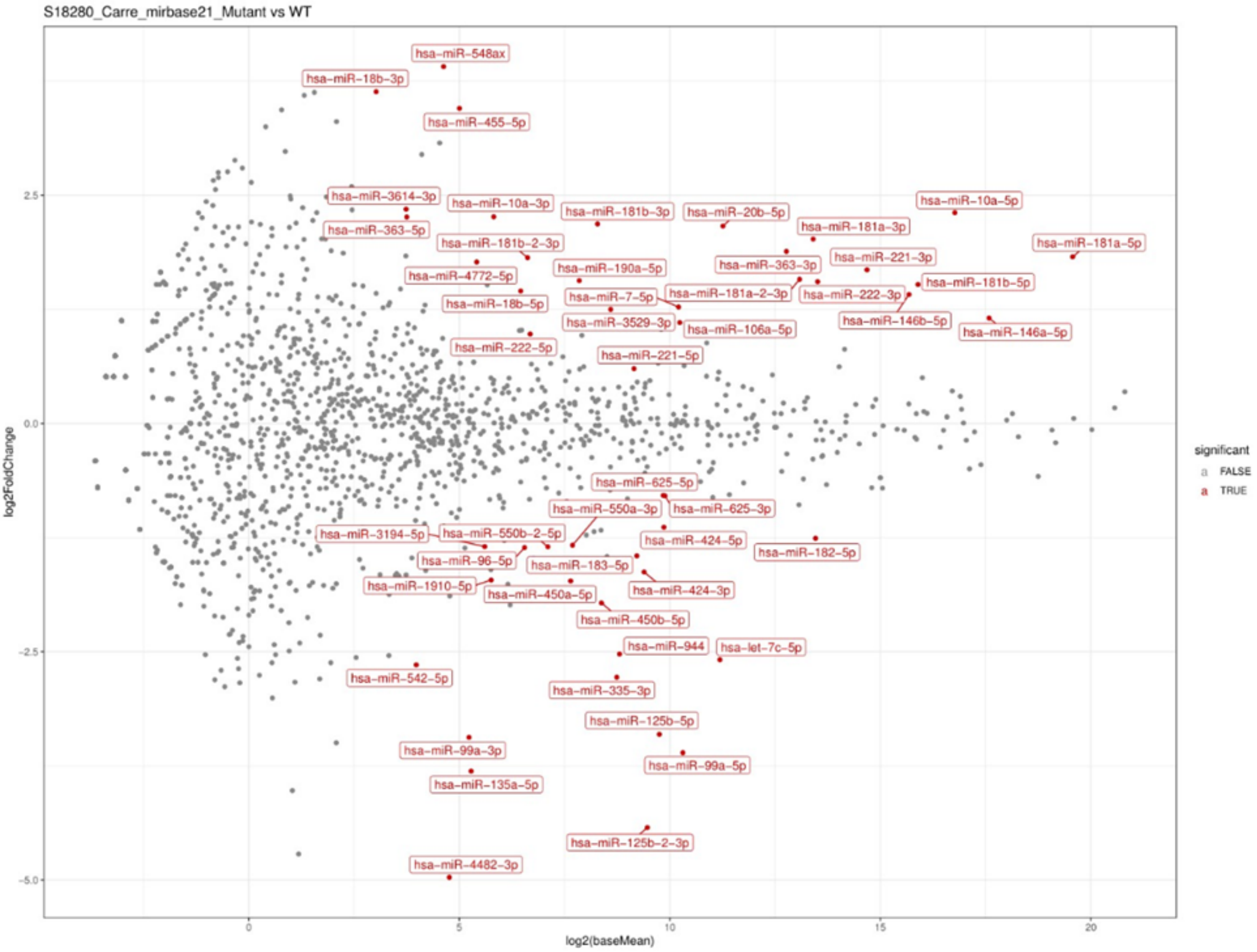


**Figure S3B. FTSJ1 loss of function leads to miRNAs deregulation in NSXLID affected individuals LCLs.** MAplot on data from sequencing of miRNAs showing multiple deregulated miRNAs (LCL WT for FTSJ1: WT; LCL Mutant for FTSJ1: Mutant).

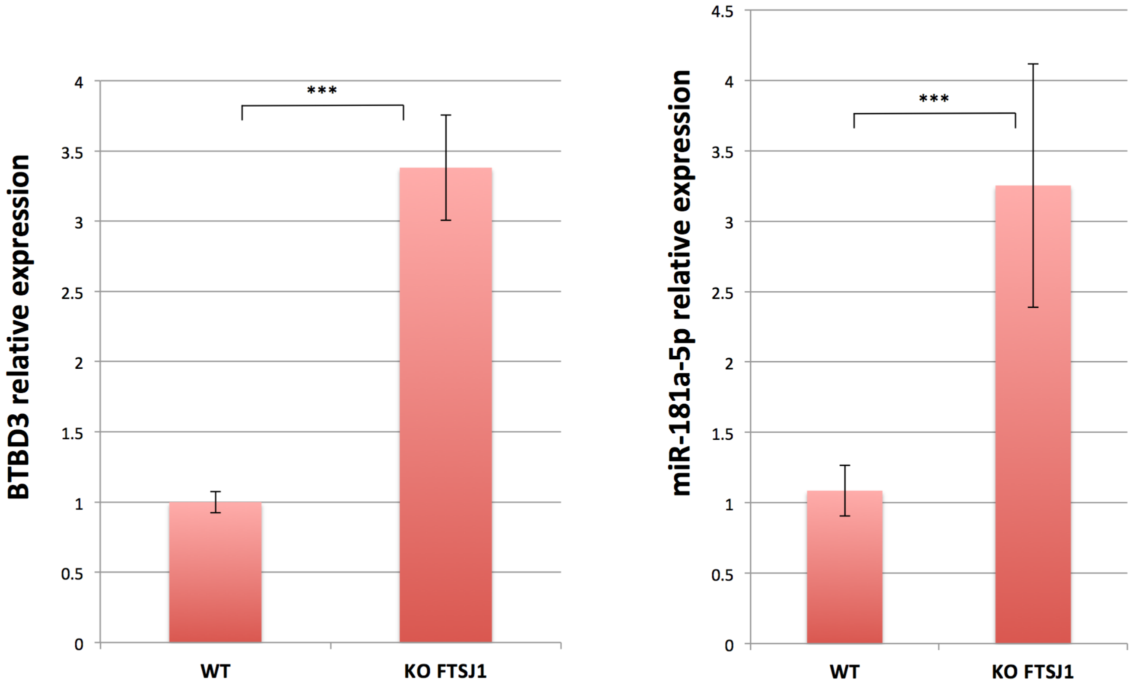


**Figure S4A. *BTBD3* and *miR-181a-5p* are expressed in HeLa cells similarly to LCLs.** RT-qPCR analysis confirms expression of *BTBD3* (Left panel) and *hsa-miR-181a-5p* (Right panel) in WT and FTSJ1 KO HeLa cells. Similarly to what is observed in patient cells, both *BTBD3* and *hsa-miR-181a-5p* are upregulated in FTSJ1 KO cells. *BTBD3* levels were normalized to *GAPDH* steady state levels, and *miR-181-a-5p* levels were normalized to the small spike-in RNA *Unisp6*. Error bars represent the standard deviation between three/six independent biological samples respectively. *BTBD3* ****p*= 4,17E-04. *miR181a-5p* *** *p*= 1,31E-04 (paired Student's t-test).

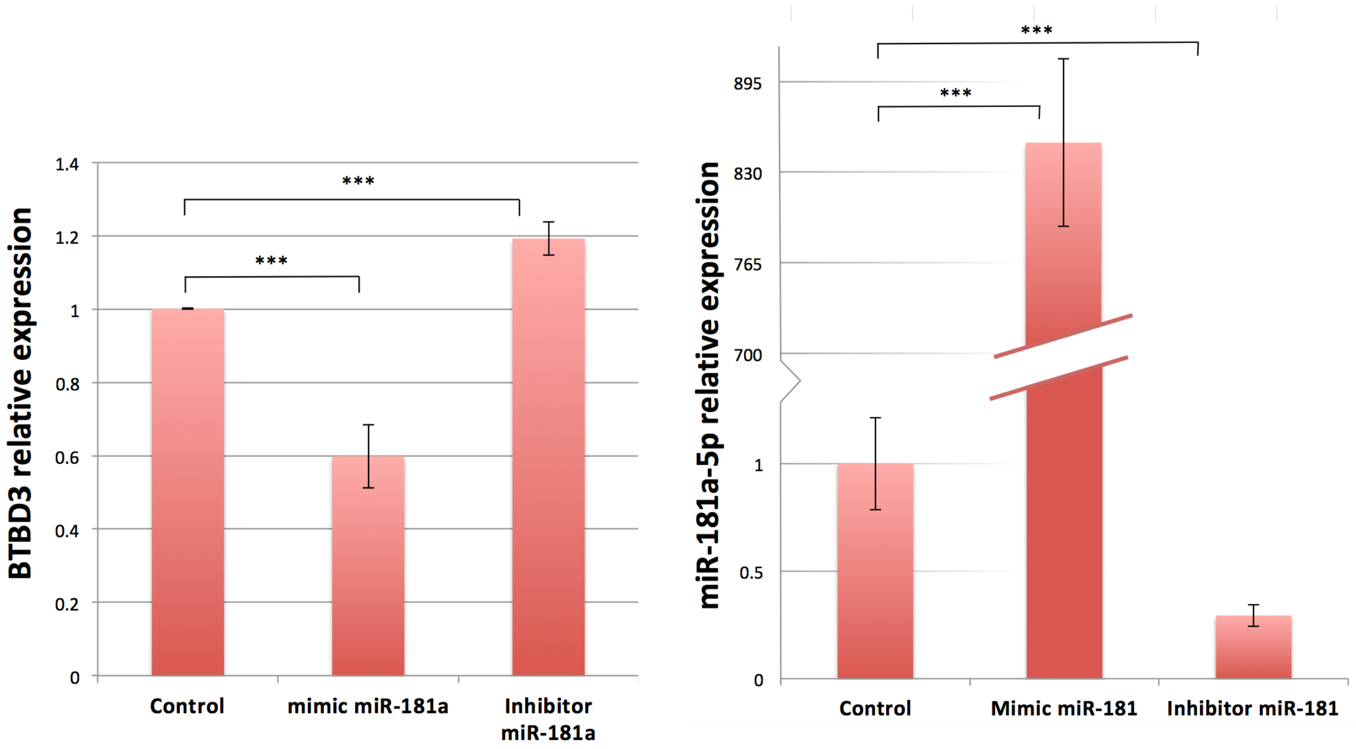


**Figure S4B. *miR-181a-5p* targets *BTBD3* in HeLa cells. Left Panel:** *miR-181a-5p* complementation in HeLa cells results in downregulation of *BTBD3* mRNA when compared to an untransfected control, as shown by relative quantification by RT-qPCR (****p*= 8,55E-05). Inhibition of *miR181a-5p* in HeLa cells results in upregulation of *BTBD3* mRNA (****p*= 1,61E-04). *BTBD3* levels were normalized to *GAPDH*. *p* values are calculated with Student's t-test on four independent biological samples. **Right Panel:** *miR-181a-5p* upregulation and downregulation is verified by RTqPCR to assess transfection efficiency. *miR-181-a-5p* levels were normalized to the small spike-in RNA *Unisp6*. Indicated *p*-values were calculated with paired Student's t-test (untransfected control/mimic *miR-181a-5p* ****p*=2,88E-04; untransfected control/inhib. *miR-181a-5p* ****p*=6,37E-04).

| # | miRNA | baseMean_mutant | baseMean_wt | log2FoldChange Mutant_vs_WT | padj |
|---|---|---|---|---|---|
| 1 | hsa-miR-20b-5p | 3987 | 538 | 2,16 | 1,21E-06 |
| 2 | hsa-miR-222-3p | 16116 | 6101 | 1,55 | 6,24E-06 |
| 3 | hsa-miR-548ax | 42 | 3 | 3,91 | 6,24E-06 |
| 4 | hsa-miR-125b-2-3p | 235 | 1295 | -4,43 | 1,37E-05 |
| 5 | hsa-miR-221-3p | 37248 | 12664 | 1,68 | 3,76E-05 |
| 6 | hsa-miR-335-3p | 217 | 690 | -2,78 | 7,92E-05 |
| 7 | hsa-miR-181b-2-3p | 142 | 44 | 1,82 | 7,92E-05 |
| 8 | hsa-miR-99a-5p | 538 | 2186 | -3,61 | 0,0001 |
| 9 | hsa-miR-10a-5p | 161396 | 49508 | 2,31 | 0,0002 |
| 10 | hsa-miR-181b-3p | 472 | 111 | 2,18 | 0,0005 |
| 11 | hsa-miR-106a-5p | 1737 | 547 | 1,10 | 0,0005 |
| 12 | hsa-miR-181a-2-3p | 12183 | 4280 | 1,58 | 0,0009 |
| 13 | hsa-miR-146a-5p | 256346 | 121011 | 1,15 | 0,0009 |
| 14 | hsa-miR-4482-3p | 2 | 58 | -4,97 | 0,0009 |
| 15 | hsa-miR-125b-5p | 468 | 1354 | -3,41 | 0,0009 |
| 16 | hsa-miR-450b-5p | 235 | 454 | -1,97 | 0,0009 |
| 17 | hsa-miR-424-3p | 608 | 750 | -1,63 | 0,0012 |
| 18 | hsa-miR-363-3p | 10299 | 2814 | 1,88 | 0,0017 |
| 19 | hsa-let-7c-5p | 1044 | 3949 | -2,59 | 0,0017 |
| 20 | hsa-miR-450a-5p | 157 | 254 | -1,73 | 0,0020 |
| 21 | hsa-miR-18b-5p | 131 | 34 | 1,45 | 0,0033 |
| 22 | hsa-miR-550a-3p | 141 | 288 | -1,33 | 0,0033 |
| 23 | hsa-miR-181a-5p | 1097695 | 379340 | 1,82 | 0,0033 |
| 24 | hsa-miR-550b-2-5p | 94 | 192 | -1,35 | 0,0044 |
| 25 | hsa-miR-181a-3p | 15949 | 4441 | 2,02 | 0,0051 |
| 26 | hsa-miR-181b-5p | 83655 | 32365 | 1,52 | 0,0100 |
| 27 | hsa-miR-183-5p | 391 | 848 | -1,45 | 0,0112 |
| 28 | hsa-miR-99a-3p | 24 | 55 | -3,44 | 0,0134 |
| 29 | hsa-miR-135a-5p | 6 | 80 | -3,81 | 0,0135 |
| 30 | hsa-miR-146b-5p | 70718 | 29704 | 1,41 | 0,0190 |
| 31 | hsa-miR-542-5p | 13 | 19 | -2,65 | 0,0321 |
| 32 | hsa-miR-944 | 138 | 833 | -2,53 | 0,0376 |
| 33 | hsa-miR-625-5p | 707 | 1186 | -0,79 | 0,0395 |
| 34 | hsa-miR-625-3p | 723 | 1220 | -0,79 | 0,0412 |
| 35 | hsa-miR-4772-5p | 56 | 26 | 1,77 | 0,0420 |
| 36 | hsa-miR-182-5p | 8091 | 15213 | -1,26 | 0,0473 |

**Table S1. FTSJ1 loss of function leads to miRNAs deregulation in NSXLID affected individuals LCLs.** A list of the significantly deregulated miRNAs and their log2 fold change and adjusted p.value between FTSJ1 loss-of-function LCLs and control LCLs.
